# Intestinal *Bacteroides* Modulates Systemic Inflammation and the Microbial Ecology in a Mouse Model of CF: Evidence for Propionate and other Short Chain Fatty Acids Reducing Systemic Inflammatory Cytokines

**DOI:** 10.1101/2022.01.05.475125

**Authors:** Courtney E. Price, Rebecca A. Valls, Alexis R. Ramsey, Nicole A. Loeven, Jane T. Jones, Kaitlyn E. Barrack, Joseph D Schwartzman, Darlene B. Royce, Robert A. Cramer, Juliette C. Madan, Benjamin D. Ross, James Bliska, George A. O’Toole

**Author notes:** To whom correspondence should be addressed Department of Microbiology and Immunology Geisel School of Medicine at Dartmouth Rm. 202 Remsen Building 66 North College Street, Hanover, NH, 03755. These authors contributed equally to this manuscript (Nicole A. Loeven, Jane T. Jones).

## Abstract

Persons with cystic fibrosis, starting in early life, have intestinal microbiome dysbiosis characterized in part by a decreased relative abundance of the genus *Bacteroides*. *Bacteroides* is a major producer of the intestinal short chain fatty acid (SCFA) propionate. We demonstrate here that CFTR-/- Caco-2 intestinal epithelial cells are responsive to the anti-inflammatory effects of propionate. Furthermore, *Bacteroides* isolates inhibit the IL-1β-induced inflammatory response of CFTR-/- Caco-2 intestinal epithelial cells and do so in a propionate-dependent manner. *Bacteroides* isolates also produce low levels of butyrate; this SCFA is positively correlated with inhibition of the inflammatory response. Finally, the introduction of *Bacteroides-*supplemented stool from infants with CF into the gut of *Cftr^F508del^* mice results in an increase in propionate in the stool as well as the reduction in several systemic pro-inflammatory cytokines. *Bacteroides* supplementation also reduced the fecal relative abundance of *E. coli*, indicating a potential interaction between these two microbes, consistent with previous clinical studies. Together, our data indicate the important role of *Bacteroides* and *Bacteroides*-derived propionate in the context of the developing microbiome in infants and children with CF, which could help explain the observed gut-lung axis in CF.

## INTRODUCTION

Cystic fibrosis (CF) is a common heritable disease with multi-organ effects that ultimately reduce quality of life and life expectancy for persons with CF (pwCF) (*Cystic Fibrosis Foundation Patient Registry 2019 Annual Data Report*, 2020). CF is caused by a defect in the cystic fibrosis transmembrane conductance regulator (CFTR), which leads to dysregulation of ion and water flow and an accumulation of thick, sticky mucus at mucosal surfaces (Riordan et al., 1989). In the lung, this mucosal alteration leads to colonization and chronic infection with polymicrobial communities of opportunistic pathogens (Bisht, Baishya, & Wakeman, 2020; Filkins & O’Toole, 2015). Physiologic effects of CFTR mutation in the gut include risk of meconium ileus at birth, longer gastrointestinal transit times, and dysbiosis of the gut microbiota (Antosca et al., 2019; De Lisle & Borowitz, 2013; Hoffman et al., 2014; Manor et al., 2016). Until relatively recently, CF microbiology research focused almost exclusively on the lung microbiota, as the majority of CF-related deaths are attributable to respiratory complications (*Cystic Fibrosis Foundation Patient Registry 2019 Annual Data Report*, 2020; Rogers et al., 2003; Surette, 2014). However, interest in the CF gut microbiome and its influence on distal organ health has grown as the role of the gut microbiome in broader health outcomes has become more apparent (Antosca et al., 2019; Burke et al., 2017; Coffey et al., 2019; Hayden et al., 2020; Hoen et al., 2015; Madan et al., 2012).

The human gut microbiome is comprised of diverse microorganisms that influence a broad range of host health outcomes, including normal immune development and function (Budden et al., 2017). Cross-talk between the gut and the lung has been referred to as the “gut-lung axis”, and this connection between gut and lung has been proposed to play a role in a number of airway diseases, including CF (Price & O’Toole, 2021). Both infants and adults with CF have a dysbiotic gut microbiome (Antosca et al., 2019; Burke et al., 2017; Fouhy et al., 2017; Hayden et al., 2020; Hoffman et al., 2014; Kristensen et al., 2020), and there is a significant correlation between gut microbiome beta-diversity for infants and children with CF and the risk of airway exacerbation in this population (Antosca et al., 2019; Hoen et al., 2015). Additionally, reduced microbial diversity in the gut microbiome correlates with earlier onset of the first clinical exacerbation in infants (Hoen et al., 2015). Studies from our group and others have demonstrated that children less than 1 year of age with CF have significantly lower relative abundance of *Bacteroides* spp. and increased relative abundance of *Escherichia coli* in their stool compared to children without CF (Antosca et al., 2019; Coffey et al., 2019; Hayden et al., 2020; Hoffman et al., 2014; Manor et al., 2016; Nielsen et al., 2016).

*Bacteroides* are primarily beneficial symbiotic microbes that promote normal immune development but can also be opportunistic pathogens (Bornet & Westermann, 2021; A. G. Wexler & Goodman, 2017). We hypothesized that a lack of *Bacteroides* may alter immune signaling in pwCF, leading to higher systemic inflammation and higher rates of clinical exacerbation. We have previously utilized an *in vitro* trans-well coculture system to demonstrate that *Bacteroides* can decrease the levels of IL-8, a pro-inflammatory cytokine, in CRISPR-modified CFTR-/CFTR- Caco-2 human intestinal epithelial cells (Antosca et al., 2019; Hao et al., 2020). Additionally, live bacteria were not required for the reduction of IL-8 levels, as supernatant from *Bacteroides* cultures also reduced the levels of this cytokine.

In this study, we utilized a larger panel of *Bacteroides* isolates and investigated the metabolomic and anti-inflammatory properties of these isolates. We found that IL-8 is broadly downregulated by many *Bacteroides* isolates and that this downregulation is dependent on the production of propionate. Furthermore, the addition of *Bacteroides* to the gut in a *Cftr^F508del^* mouse model of CF demonstrated that the presence of intestinal *Bacteroides* can impact systemic inflammation as well as shift the microbial populations of the fecal microbiota. This work demonstrates that *Bacteroides* may be a key driver of the gut-lung axis in CF.

## RESULTS

### Supernatants from *Bacteroides* Isolates Downregulate IL-8

Previous work from our group utilized a CFTR-/- Caco-2 intestinal epithelial cell line developed by Hao et al. (Hao et al., 2020) to demonstrate that *Bacteroides* supernatants can downregulate the TNF-*α*-stimulated production of IL-8 in a trans-well coculture system (Antosca et al., 2019). In order to increase experimental throughput, we chose to work with cells cultured on plastic in 24 and 96-well plates. Previous studies demonstrated that Caco-2 intestinal epithelial cells cultured on plastic are more responsive to IL-1β stimulation than TNF-*α* or LPS (Schuerer-Maly, Eckmann, Kagnoff, Falco, & Maly, 1994). We therefore tested the response of three CFTR-/- and three WT Caco-2 intestinal epithelial cell lines cultured on plastic to stimulation with 10ng/mL IL- 1β for 24 hours. IL-1β stimulated robust production of IL-8 in all cell lines (Fig. S1A-B), but with considerable variability dependent on both cell line and experimental replicate. All subsequent experiments were performed in the CFTR-/- S1 cell line as this line was used in our previous work (Antosca et al., 2019) and showed a relatively consistent response in terms of IL-8 production when treated with IL-1β. A 24-hour time course of IL-1β stimulation of CFTR-/- S1 cells demonstrated that IL-8 accumulates rapidly over the first 6 hours of IL-1β exposure and then more slowly up to 24 hours (Fig. S1C).

Next, we tested the impact of supernatants derived from *Bacteroides* and *Parabacteroides* isolates on the IL-1β-stimulated IL-8 production by CFTR-/- S1 Caco-2 intestinal epithelial cells (hereafter called “CFTR-/- Caco-2 cells”). We isolated these *Bacteroides* and *Parabacteroides* strains from the stool of children with and without CF (Table 1). Strains with the prefix CFPLTA were isolated from healthy children, while all other clinical isolates were from the stool of children with CF. Bacterial supernatants were prepared from 19 isolates in supplemented MEM (sMEM, see Materials and Methods) and screened *in vitro* for their effects on the inflammatory response of IL-1β- stimulated CFTR-/- Caco-2 cells (Fig. S2A). The majority of the *Bacteroides* isolates trended towards downregulation of IL-8, while supernatants from *Parabacteroides* tended to have no effect or to increase IL-8 production. Supernatant treatment did not significantly decrease the viability of the CFTR-/- Caco-2 cells, but treatment with *Parabacteroides* supernatant trended towards increased viability relative to IL-1β alone (Fig. S2B). Growth of *Bacteroides* and *Parabacteroides* varied in sMEM in an isolate-dependent manner (Fig. S2C). In general, *Bacteroides* grew better than *Parabacteroides* in sMEM, which may partially explain the stronger down-regulatory effects on IL-8 by *Bacteroides* isolates.

**Table 1.** List of all bacterial isolates.

| Strain ID | Genus/Species | Source | SMC ID | Assembly |
| --- | --- | --- | --- | --- |
| AD126T_3B | *Bacteroides fragilis* | CF | 9092 |  |
| AD135F_3B | *Bacteroides fragilis* | CF | 9093 | GCA_007896605.<br>1 |
| HAP130N_2B | *Bacteroides fragilis* | CF | 9094 |  |
| TL139C_3B | *Bacteroides fragilis* | CF | 9095 |  |
| CFPLTA004_1B | *Bacteroides fragilis* | non-CF | 9096 | GCA_007896595.<br>1 |
| CFPLTA007_1B | *Bacteroides fragilis* | non-CF | 9097 |  |
| ATCC_29741 | *Bacteroides thetaiotaomicron* | non-CF | 7758 |  |
| AD135X_1B | *Bacteroides thetaiotaomicron* | CF | 7764 | GCA_009025475.<br>1 |
| VPI (LAB STRAIN) | *Bacteroides thetaiotaomicron* | CF | 7757 |  |
| ANK132K_1B | *Bacteroides vulgatus* | CF | 9098 | GCA_007896765.<br>1 |
| CFPLTA002_1B | *Bacteroides vulgatus* | non-CF | 9099 | GCA_009025805.<br>1 |
| CFPLTA003_2B | *Bacteroides vulgatus* | non-CF | 9100 | GCA_007896665.<br>1 |
| RH127O | *Bacteroides vulgatus* | CF | 9101 | GCA_009025535.<br>1 |
| TL139H | *Bacteroides vulgatus* | CF | 9102 |  |
| ADR1290_2B | *Parabacteroides distasonis* | CF | 9103 | GCA_009024595.<br>1 |
| CFPLTA001_1B | *Parabacteroides distasonis* | non-CF | 9104 |  |
| CFPLTA003_1B | *Parabacteroides distasonis* | non-CF | 9105 | GCA_007896695.<br>1 |
| CFPLTA006_1B | *Parabacteroides distasonis* | non-CF | 9106 | GCA_009025485.<br>1 |
| AD135X_2B | *Parabacteroides merdae* | CF | 7765 |  |

*Bacteroides* from pwCF did not have significantly different impacts on IL-8 compared with *Bacteroides* isolates from healthy children (Fig. S3A). We found a modest negative correlation between *Bacteroides* growth and IL-8 production by simple linear regression (R^2^ = 0.075, p = 0.039), as well as a positive correlation between IL-8 production and CFTR-/- Caco-2 cellular viability (R^2^ = 0.103, p = 0.018; Fig. S3B-C). In summary, IL-8 downregulation is primarily dependent on isolate, is modestly influenced by CFTR-/- Caco-2 cellular viability and *Bacteroides* growth, and is independent of isolate origin.

A subset of *Bacteroides* isolates were selected, supernatants from these strains were prepared, and re-tested for their impact on IL-8 production by IL-1β-stimulated CFTR-/- Caco-2 cells grown in 24-well plates. A control with no IL-1β was included, showing that baseline IL-8 production is low (∼10 ng/uL; Fig. 1A). IL-8 secreted from the CFTR-/- Caco-2 cells was measured by ELISA after exposure to bacterial supernatants for 6 or 24 hours (Figure 1A-B). We identified two isolates, *B. vulgatus* CFPLTA003_2B and *B. fragilis* AD126T_3B, that had consistent, strong IL-8 downregulatory effects after 6 hours of coculture (Fig. 1A). The downregulatory effects of *Bacteroides* supernatants were reduced after 24 hours, but IL-8 still trended downwards in the presence of supernatants (Fig. 1B). Overall, *Bacteroides* supernatants inhibit the production of IL-8 by CFTR-/- Caco-2 cells stimulated with IL- 1β.

**Figure 1.**
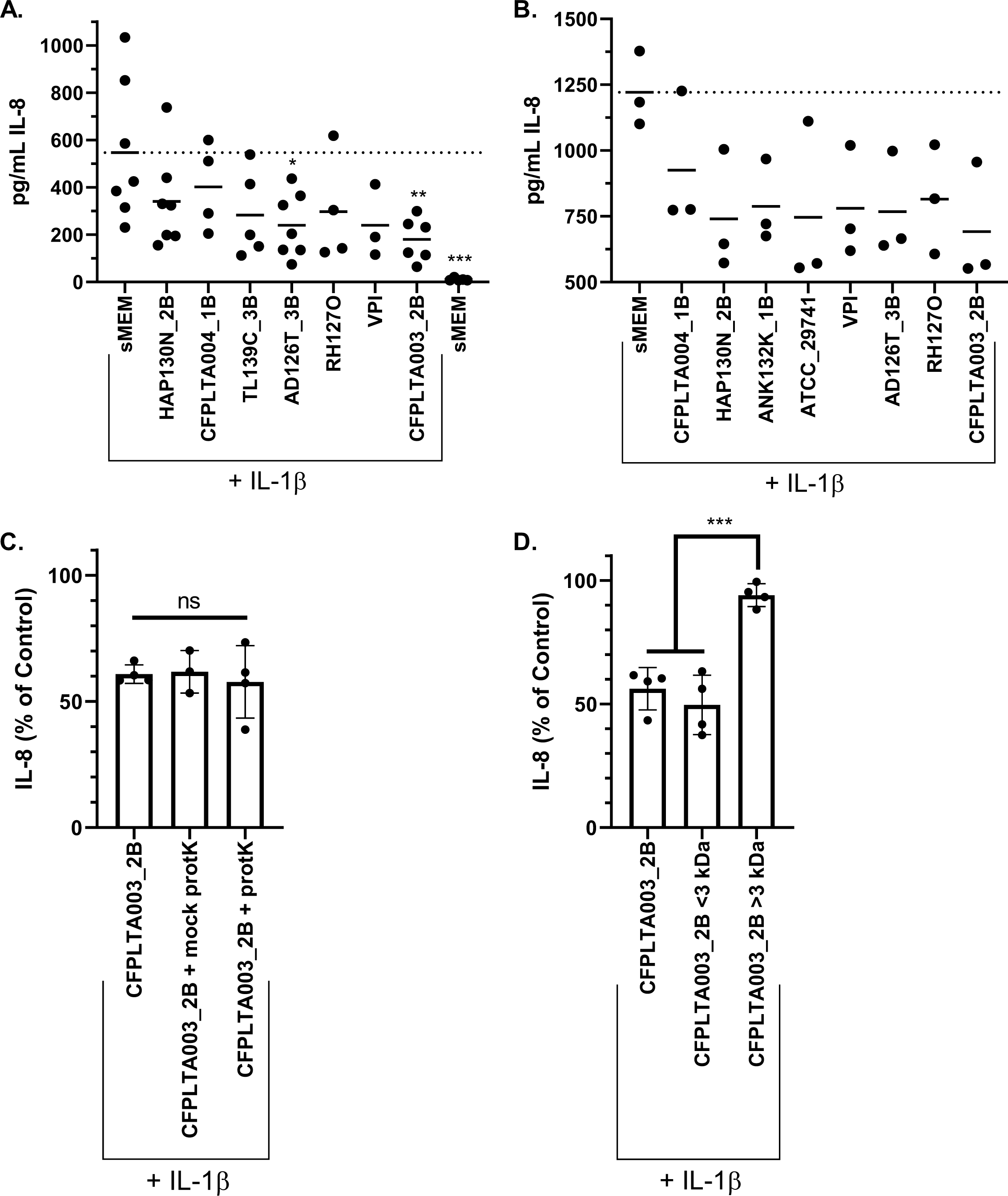
*Bacteroides* supernatants downregulate IL-8 production by Caco-2 cells and IL-8 downregulation is likely not due to a protein. CFTR-/- Caco-2 cells were cultured for 2 weeks in 24-well plates and then exposed to *Bacteroides* supernatants prepared in sMEM as described in the Materials and Methods. Each point represents the average of 4-6 technical replicates from a single biological replicate. IL-8 was quantified by ELISA after A) 6 hours and B) 24 hours of CFTR-/- Caco-2 cell’s exposure to the bacterial supernatants from the indicated *Bacteroides* strains. Significance was determined by unpaired one-way ANOVA followed by Dunnett’s post-test with MEM + IL-1β as the reference. *p<0.05, **p<0.01, ****p<0.0001. C-D) IL-8 was quantified by ELISA after 24 hours of CFTR-/- Caco-2 cells exposure to the bacterial supernatants from the indicated treatment. IL-8 quantity is displayed as a percentage of the MEM + IL-1β control in the same biological replicate. Effects of C) proteinase K treatment and D) size fractionation on IL-8-reducing activity. Significance was tested by unpaired one-way ANOVA followed by Tukey’s post-test. **p< 0.01.

### The Majority of Our *Bacteroides* Isolates Do Not Encode *Bacteroides fragilis* Toxin

Enterotoxigenic strains of *Bacteroides fragilis* produce a protein known as *Bacteroides fragilis* enterotoxin (BFT) that is implicated in many intestinal inflammatory disorders, including development of colorectal cancer (Valguarnera & Wardenburg, 2020; H. M. Wexler, 2007). BFT is located on a mobilizable pathogenicity island and thus has the potential to be transferred between isolates (Franco et al., 1999; Moncrief, Duncan, Wright, Barroso, & Wilkins, 1998). Furthermore, BFT can induce IL-8 (Wu et al., 2004), which may interfere with IL-8-downregulating factors in our assay. To rule out the possibility that BFT was interfering with IL-8 down regulation, we probed the genomes of our *B. fragilis* isolates for their potential to encode this toxin. Whole genome sequencing data is available for a subset of the isolates in this study (PRJNA557692). We used BLAST to assess all proteins from each isolate for BFT (See Materials and Methods for details). Significant sequence similarity to BFT was identified in the isolate TL139C_3B. Interestingly, this isolate moderately downregulates IL-8 (Fig. S2A-B), indicating that if BFT is expressed under our experimental conditions, it does not eliminate IL-8 downregulation by this strain.

### Effects of *Bacteroides* Supernatants on IL-8 Production is Due to a Heat-Stable, Small Molecular Weight Factor

We selected a single *B. vulgatus* isolate, CFPLTA003_2B, to work with for mechanistic studies; this isolate was chosen for its consistent and strong downregulation of IL-8. We treated supernatants from *B. vulgatus* isolate CFPLTA003_2B with proteinase K (protK) to determine whether *Bacteroides* reduction of IL-8 was protein-dependent (see Materials and Methods for details). Supernatant was treated with protK for 1-2 hours at 37°C, and protK was subsequently inactivated by heating at 95°C for 15 minutes in a sealed plastic tube. A mock condition, where supernatant underwent the same temperature incubation steps as the protK-treated samples was also included. Supernatant from *B. vulgatus* CFPLTA003_2B downregulated IL-8 from standard, mock, and protK-treated conditions (Fig. 1C). The lack of sensitivity to protK and extended heat treatment indicated that a protein is likely not required for supernatant-mediated IL-8 downregulation and that the factor(s) required for this activity is heat stable.

We next used a 3 kDa size cut-off filter to estimate the size of the factor(s) responsible for IL-8 downregulation. *B. vulgatus* CFPLTA003_2B supernatant was placed in a 3 kDa spin-column, and CFTR-/- Caco-2 cells were incubated with either the flowthrough containing small molecular weight molecules (<3 kDa) or the retentate fraction containing molecules >3 kDa, which was subsequently diluted to its original volume. The flowthrough, but not the retentate fraction reduced IL-8 production (Fig. 1D), supporting the conclusion that a small molecular weight molecule was responsible for this activity.

### IL-8 production by the CFTR-/- Caco-2 Cell Line is Responsive to SCFAs

Short chain fatty acids (SCFAs) are generated by microbial fermentation in the intestine and are an important aspect of the microbiome that is beneficial to host health, as SCFAs both provide an important source of energy for colonic epithelial cells and regulate inflammatory host responses (Parada Venegas et al., 2019). The three highest-abundance SCFAs in the intestine are acetate, propionate, and butyrate (Parada Venegas et al., 2019). *Bacteroides* have been reported to produce the two SCFAs propionate and acetate, but not butyrate (Louis & Flint, 2017). We hypothesized that *Bacteroides*-derived propionate or acetate might be reducing IL-8 in our coculture experiments (see Fig. 1). We tested the dose-responsiveness of these three abundant SCFAs and found that CFTR-/- Caco-2 cells respond to all three, albeit at different concentrations (Fig. 2).

**Figure 2.**
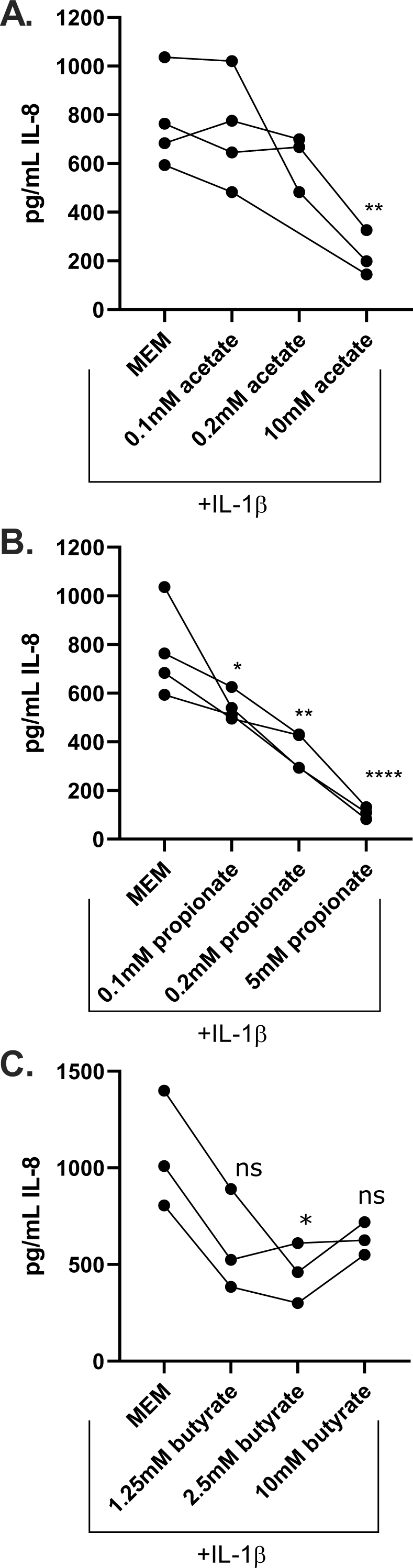
CFTR-/- Caco-2 cells are responsive to anti-inflammatory effects of SCFAs. CFTR-/- Caco-2 cells were cultured for 2 weeks in 24-well plates and then treated with IL-1β alone, or with the addition of the indicated concentrations of SCFAs. IL-8 was quantified by ELISA after 24 hours of culture with A) sodium acetate B) sodium propionate or C) sodium butyrate at the indicated concentration. Each point indicates the average of 4-6 technical replicates from a single biological replicate, with 3-4 independent biological replicates performed. Lines connect results from experiments performed on the same day. Significance was tested by unpaired one-way ANOVA followed by Dunnett’s post-test with MEM + IL-1β as the reference. *p<0.05, **p< 0.01, ****p<0.0001.

The CFTR-/- Caco-2 cells are responsive to acetate starting at 10mM (Fig. 2A) but are very responsive to propionate at a concentration as low as 0.1mM (Fig. 2B). Neither acetate nor propionate impact cellular viability at the concentrations tested (Fig. S4A-B). The IL-8 inflammatory response is not linearly responsive to butyrate; downregulation occurs at 2.5mM but not at 10mM; this observation is likely because butyrate begins to inhibit cellular viability at a 10mM concentration (Fig. S4C). In summary, CFTR-/- Caco-2 cells are responsive to the anti-inflammatory effects of major intestinal SCFAs.

### A Strain Deficient in Propionate Synthesis Reduces Anti-Inflammatory Properties of *B. thetaiotaomicron*

We utilized *B. thetaiotaomicron* BT1686-89, a mutant lacking the methyl malonyl-CoA transcarboxylase gene, which is required for the conversion of succinate to propionate and is deficient in propionate production, to test whether propionate is necessary for downregulation of IL-8 observed for *Bacteroides* supernatants (Kovatcheva-Datchary et al., 2015). BT1681-89 was constructed in the *B. thetaiotaomicron* VPI-5482 Δ*tdk* background and is referred to as *B. thetaiotaomicron Δprp* in this report. The wild-type *B. thetaiotaomicron* VPI-5482 and *B. thetaiotaomicron Δtdk* isogenic parents are both included as controls in these studies. Supernatants were prepared from each strain in sMEM and the supernatants applied to the CFTR-/- Caco-2 cells for 6 and 24 hours as described in the Materials and Methods and in Figure 3.

**Figure 3.**
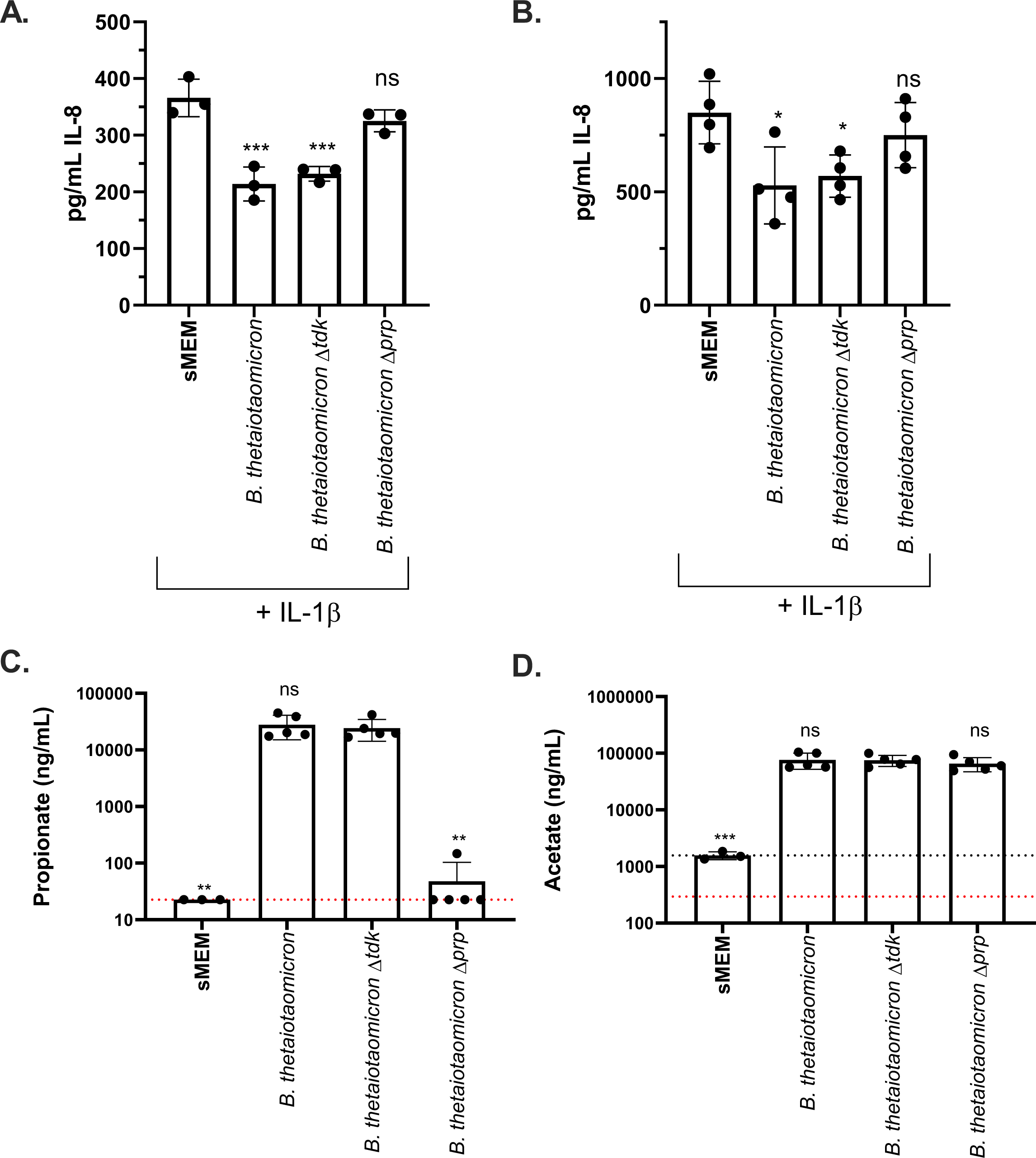
IL-8 downregulation by *B. thetaiotaomicron* is propionate dependent. CFTR-/- Caco-2 cells were cultured for 2 weeks in 24-well plates and then cocultured with *B. thetaiotaomicron* supernatants from the indicated strains prepared in sMEM as described in the Materials and Methods. IL-8 was quantified by ELISA after A) 6 hours and B) 24 hours of the CFTR-/- Caco-2 cell’s exposure to the bacterial supernatants from the indicated strains. Each point represents the average of 4-6 technical replicates from a single biological replicate, with 3 independent biological replicates performed. Lines connect data from the same experiment. Significance was determined by unpaired one-way ANOVA followed by Dunnett’s post-test with sMEM + IL-1βas the reference. *p<0.05,**p<0.01, ****p<0.0001. The SCFAs propionate (panel C) and acetate (panel D) were quantified by GC-MS. The dashed red line indicates the assay limit of detection (LOD). In panel D the dashed black line indicates the average acetate concentration detected in the medium, as this medium contains acetate above the LOD. Significance was determined by unpaired one-way ANOVA followed by Dunnett’s post-test with *B. thetaiotaomicron Δtdk* as the reference. **p<0.01.

While supernatants from both wild-type *B. thetaiotaomicron* and *B. thetaiotaomicron Δtdk* strains significantly downregulated IL-8 production at 6 and 24 hours, the *B. thetaiotaomicron Δprp* did not show such activity (Fig. 3A-B). Each bacterial culture was also plated for CFU/mL and there was no significant difference between any of the three strains (Fig. S5A), indicating that differential growth of the strains was not responsible for the observed impact on IL-8 production. XTT assays were performed for each experiment to assess the viability of the CFTR-/- Caco-2 cells after treatment with the supernatants; there was no difference in viability caused by treatment with the supernatants from the three strains (Fig. S5B-C).

SCFAs were also quantified from supernatants from each strain by GC-MS, confirming that *B. thetaiotaomicron Δprp* did not produce propionate in our medium, and that the quantity of other measured SCFAs was not altered by the loss of the methyl malonyl-CoA transcarboxylase gene and the resulting loss of propionate production (Fig. 3C-D, Fig. S6). Together, these results demonstrate that *Bacteroides* downregulation of IL-8 is dependent on propionate production.

### *Bacteroides* Isolates Produce Different Quantities of a Subset of SCFAs

We next wanted to determine whether there were strain-dependent differences in SCFA production by *Bacteroides* isolates, and whether these differences would correlate with the ability of isolates to downregulate IL-8. We therefore prepared supernatants from the 19 isolates that we originally screened for IL-8 downregulation (Fig. S2) and quantified SCFAs in each supernatant by GC-MS. Unsurprisingly, acetate and propionate were the primary metabolites produced by all isolates of *Bacteroides* and *Parabacteroides* (Fig. 4A, Fig. S7A-B). Other detectable SCFAs at lower concentrations included 2-methyl propionate, butyrate, 3-methyl butyrate, hexanoate and heptanoate. Pentanoate and 2-methyl pentanoate were not present above the limit of detection.

**Figure 4.**
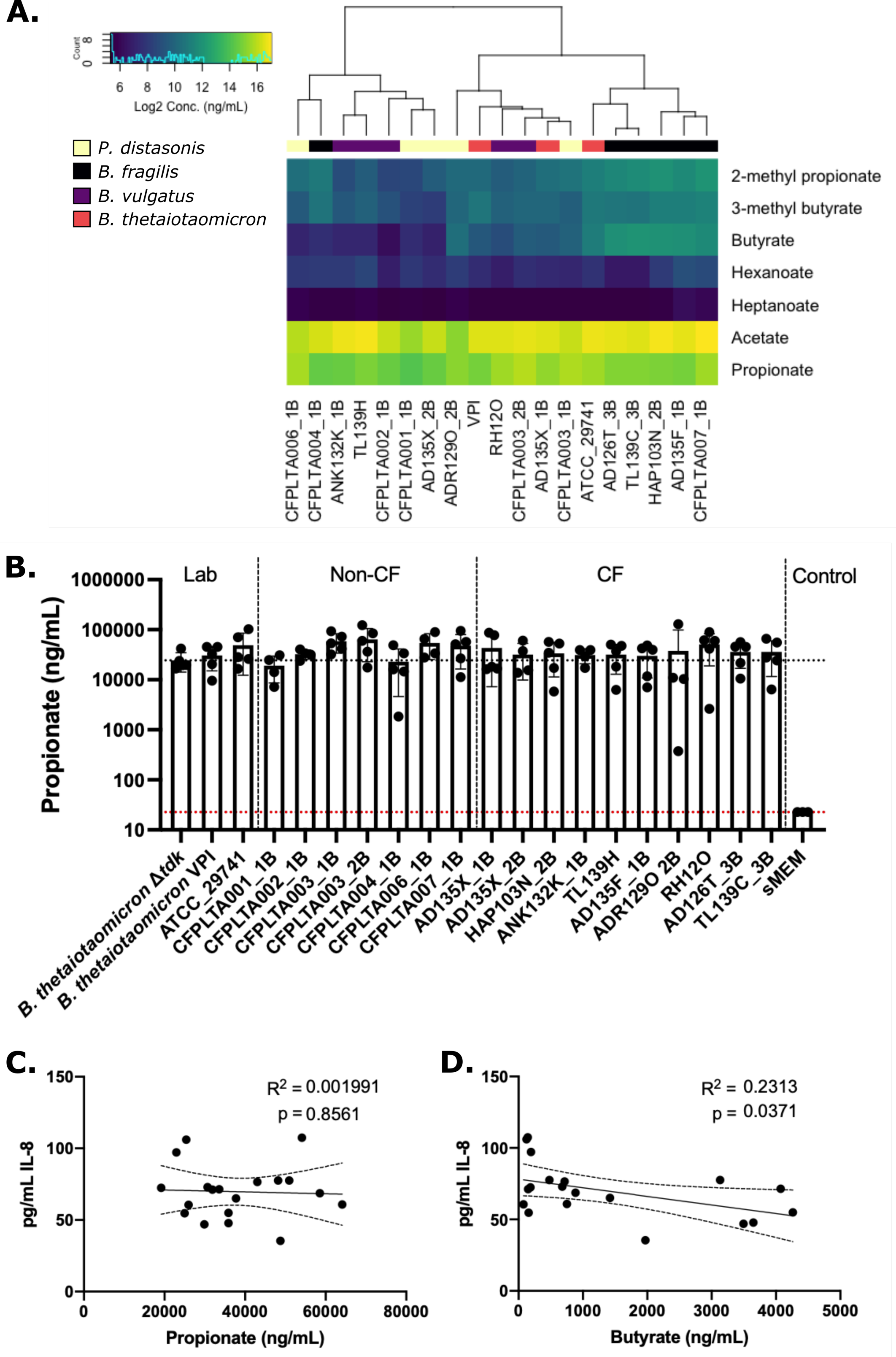
Butyrate correlates with IL-8 downregulation. SCFAs were quantified in undiluted, sterile-filtered supernatants by GC-MS as described in the Materials and Methods. A) Heatmap of Log2 transformed concentration (ng/mL) of each detectable SCFA. Pentanoic and 2-methyl pentanoates were not detected above the LOD. B) Propionate produced by individual isolates. The dashed red line indicates the LOD. The dashed black line indicates the average propionate produced by control strain *B. thetaiotaomicron Δtdk.* Linear regression was performed and is displayed with 95% confidence intervals for C) propionic and D) butyrate versus the % reduction in IL-8 level. Inset: Simple linear regression was used to calculate R^2^ and p value indicated.

Clustering of SCFAs only demonstrated partial separation by species (Fig. 4A). We did detect some species-specific differences in butyrate, 2-methyl propionate, and 3-methyl butyrate, all of which were significantly higher in *B. fragilis* isolates (Fig. S7C- E). Furthermore, acetate was significantly lower in supernatant from *Parabacteroides* isolates compared to *Bacteroides* isolates (Fig. S7A). At the isolate level, the *B. vulgatus* CFPLTA003_2B isolate produced the highest concentration of propionate (Fig. 5B; Fig. S7B). However, this difference was not statistically significantly higher than other supernatants, suggesting that propionate is necessary but not sufficient to explain the robust anti-inflammatory properties of CFPLTA003_2B, a point we address further below.

**Figure 5.**
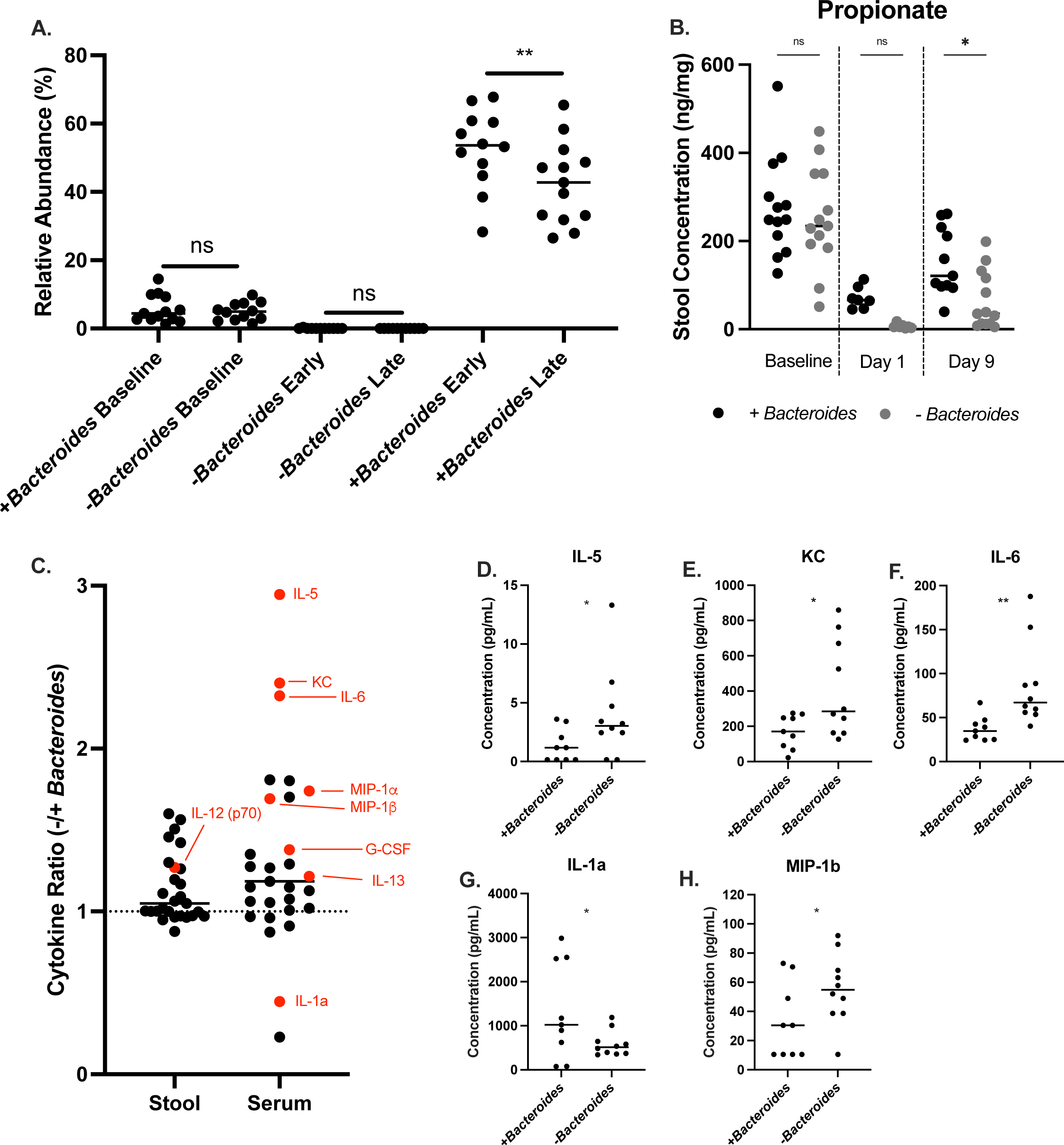
A *Cftr^F508del^* mouse model reveals that intestinal supplementation of *Bacteroides* reduces systemic cytokine levels. A-C) Stool samples were collected from *Cftr^F508del^* mice either before the onset of antibiotic treatment (Baseline) or post- gavage with pooled CF stool +/- *Bacteroides.* Each point represents data from a single mouse stool in all graphs. A) Microbial composition of stool was profiled for each mouse by 16S amplicon sequencing to determine *Bacteroides* relative abundance. ‘Early’ indicates that the stool was collected 2-4 days post-gavage and ‘Late’ indicates collection 12-13 days post-gavage. B) Propionate was quantified by GC-MS from mouse stool samples on the indicated days. A-B) Statistical significance was analyzed by one-way ANOVA with Dunnett’s post-test. *p<0.05. **p<0.01. C) Serum was collected at the end of the experiment after LPS challenge and stool was collected on the final day of sampling before LPS challenge. Cytokines were quantified by Luminex 32-plex panel for the indicated cytokines. Cytokines are displayed as a ratio of the average cytokine concentration in the (-) *Bacteroides* condition divided by the average cytokine concentration in the (+) *Bacteroides* condition. Statistical significance was analyzed by linear model and cytokines with p < 0.1 are labeled in red. D-H) Serum cytokines quantified by Luminex that were statistically significant between the (+) and (-) *Bacteroides* conditions by linear model. *p<0.05. **p<0.01. D) IL-5 (Interleukin 5), E) KC (CXCL1; IL-8 homolog) F) IL-6 (Interleukin 6), G) IL-1*α* (Interleukin 1*α*), and H) MIP- 1β (Macrophage inflammatory protein 1β).

For each SCFA detected, we used linear regression of SCFA quantity versus IL- 8 to test whether additional *Bacteroides*-derived SCFAs might impact the inflammatory response. Propionate concentrations were not significantly correlated with IL-8 levels (Fig. 5C). Interestingly, the only SCFA significantly correlated with IL-8 downregulation was butyrate (Fig. 5D). Current literature does not describe *Bacteroides* as a butyrate-producer (Louis & Flint, 2017). However, we measured low but detectable concentrations of butyrate for every *Bacteroides* and *Parabacteroides* sample tested (Fig. S7D), and the observation that butyrate concentration correlates with IL-8 production suggests production of this SCFA has biologically-relevant impacts on host cells. These results demonstrate that *Bacteroides* is dependent on multiple metabolites to reduce IL-8 production, likely including propionate and butyrate, and that low levels of *Bacteroides*-derived butyrate production may contribute to this downregulation.

### *Bacteroides* Species-Level Metabolite Signatures

The identification of multiple SCFAs of interest raised the question of the role of other metabolites produced by *Bacteroides* might impact the regulation of the inflammatory response by intestinal epithelia. We used LC-MS to generate metabolomic profiles of several *Bacteroides* isolates cultured in MEM for 48 hours or cultured in sMEM for 24 hours. Clustering of metabolite profiles demonstrated that *Bacteroides* produced significantly more metabolites in MEM versus sMEM, likely due to increased culture time (48 hrs vs. 24 hrs, respectively) under these conditions (Fig. S8A). Metabolites were measured for four isolates in MEM, and individual biological replicates clustered well at the isolate and species level (Fig. S8B). PCA plots of all biological replicates demonstrated clustering by isolate, with the two *B. vulgatus* isolates CFPLTA003_2B and RH127O clustering more closely than the two *B. thetaiotaomicron* isolates (Fig. S8C).

LC-MS analysis of the metabolites produced by isolates cultured in sMEM demonstrated higher day-to-day variability, with individual biological replicates not clustering well together (Fig. S9A). However, a PCA plot of all metabolites detected demonstrated that there was some species-level separation, with *B. thetaiotaomicron* separating from *B. fragilis* and *B. vulgatus* (Fig. S9B).

We identified twenty-four metabolites with at least a 3-fold increase in every supernatant over the medium control (Fig. S10A). The three metabolites with the highest average relative abundance increase were lactate, malate, and succinate. We tested each of these metabolites alone and found that lactate modestly down-regulated IL-8 secretion at concentrations of 5mM or greater, while neither succinate nor malate down-regulated IL-8. Indeed, malate enhanced IL-8 production at high concentrations (Fig. S10B).

We next screened a panel of a total of 26 commercially available metabolites identified by LC-MS in the *Bacteroides* supernatant (Fig. S10C). Most of the metabolites had no effect or increased IL-8 production, or if they did reduce IL-8 production this impact was due to their cytotoxic effects versus the CFTR-/- Caco-2 cells (Fig. S10C). Only one compound of the 26 tested, riboflavin, showed a large reduction of IL-8 production with no detectable cytotoxicity versus the CFTR-/- Caco-2 cells (Fig. S10C). Thus, future studies are warranted to address the impact of riboflavin to modulate IL-8 production at physiological concentrations.

### Supplementing *Bacteroides* Isolates Alters SCFAs and Cytokine Production in a *Cftr^F508del^* Mouse Model

To test the impact of the addition of *Bacteroides* to the CF gut microbiome, we designed a mouse experiment wherein the native gut microbiome of specific pathogen- free *Cftr^F508del^* mice was first suppressed by sustained antibiotic treatment and then replaced by oral gavage with a stool pool from children with CF. The stool pool contained either three *Bacteroides* isolates *B. vulgatus* CFPLTA003_2B, *B. thetaiotaomicron* SMC7758, and *B. fragilis* AD126T 3B (Table 1) or PBS only as a control, such that the two conditions were mice with a CF-like gut microbiota with and without the presence of added *Bacteroides* strains. Mouse stool was collected prior to the beginning of antibiotic treatment and for two weeks post-gavage for analysis of 16S rRNA gene-based microbiome structure, cytokines, IgA, and SCFA content (Fig. S11A). Two weeks after introducing the stool pools, mice were intranasally challenged with 0.2mg/kg *P. aeruginosa* LPS and then sacrificed 24 hours post-challenge. At the time of sacrificing, serum was collected for analysis of cytokines and SCFAs. Additionally, lung and intestinal tissues were collected for analysis of the inflammatory response via H&E staining.

Analysis of the 16S rRNA gene data across all samples showed that the average *Bacteroides* relative abundance was 5.4% at baseline (that is, before any treatments; Fig. S11B). CF stool pools used to gavage mice for the (+) and (-) *Bacteroides* groups contained a mean relative abundance of 83.4% and 0.33% *Bacteroides*, respectively (Fig. S11B). Post-gavage *Bacteroides* relative abundance was 0.02% for the (-) *Bacteroides* condition, and 47.4% for the (+) *Bacteroides* condition (Fig. S11B). Samples collected for amplicon sequencing within 2-4 days post-gavage were grouped as ‘early’ and those collected on day 12 or 13 post-gavage were grouped as ‘late’. *Bacteroides* relative abundance did not significantly change between early and late samples in the (-) *Bacteroides* condition (Fig. 5A). *Bacteroides* relative abundance remained significantly higher than baseline throughout the post-gavage period but did decrease from an average relative abundance of 53.6% early to 42.8% late in the (+) *Bacteroides* condition (Fig. 5A).

Overall community structure of these samples was analyzed by multidimensional scaling (MDS) ordination based on Bray-Curtis dissimilarity coefficients (Fig. S11C).

The overall composition of the (+) and (-) *Bacteroides* conditions were significantly different by Permanova (p = 0.014) 2-4 days post-gavage but only marginally significantly different (p = 0.085) 12-13 days post-gavage. The community was further analyzed at both time points at the genus level for significantly altered relative abundance of ASVs by DESeq2 (Fig. S11E-F). Interestingly, few ASVs were significantly different between the two conditions and only three were altered in the same direction at both time points. Unsurprisingly, the relative abundance of *Bacteroides* was increased in the (+) *Bacteroides* condition at both time points. Both *Parabacteroides* and *Escherichia* were also reduced in the (+) *Bacteroides* relative to the (-) *Bacteroides* condition at both time points. These data show that the *Bacteroides* spp. introduced by gavage were effectively retained during the course of the experiment and may influence the abundance of key CF-associated intestinal microbes, specifically *E. coli*. However, introduction of *Bacteroides* did not radically alter the overall stool microbiota composition, particularly at the later time points.

The presence of *Bacteroides* significantly increased the amount of detectable propionate and 2-methyl propionate but not acetate or butyrate in the stool (Fig. 5B; Fig. S12). However, the amount of detectable butyrate trended upwards in the (+) *Bacteroides* condition and both 2-methyl propionate and 3-methyl butyrate were significantly increased one day post-gavage (Fig. S12).

Next, to assess whether the presence of *Bacteroides* and subsequent changes in local SCFA concentrations impacted either local or systemic inflammation, we quantified both stool and serum cytokines using a Luminex 32-plex cytokine panel (Fig. 5C-H; Fig. S13-S14). Significant changes between conditions were tested by linear model and the average ratio of cytokines in the (-) *Bacteroides* condition divided by the (+) *Bacteroides* condition was graphed for each cytokine (Fig. 6C). Interestingly, 19 of the 29 stool cytokines detected trended higher in the (-) *Bacteroides* condition, but only one cytokine, IL-12 (p70), was significantly increased in the (-) *Bacteroides* condition and no cytokines were significantly increased in the (+) *Bacteroides* condition.

The majority of serum cytokines followed a similar pattern, with 23 of the 29 cytokines detected trending higher in the (-) *Bacteroides* condition. Five pro-inflammatory serum cytokines were significantly altered in serum, four of which (KC (CXCL1), IL-5, IL-6, and MIP-1b) were increased in the in the (-) *Bacteroides* condition while IL-1*α* was decreased in the (-) *Bacteroides* condition (Fig. 6D-H). Three additional serum pro-inflammatory cytokines (G-CSF, IL-13, MIP-1*α*) were also marginally significantly increased in the (-) *Bacteroides* condition (Fig 6C; Fig. S14).

Local and systemic inflammation was also assessed by quantification of calprotectin and IgA in the stool and analysis of H&E stained lung and intestinal tissue (Fig. S15-S16; see supplemental figure legend for details), but no significant differences were detected between conditions.

To assess whether increased local SCFAs corresponded with an increase in systemic SCFAs, we also quantified 12 SCFAs in the serum collected at the time of sacrificing. Only 3-methyl valerate, which was modestly higher in the (+) *Bacteroides* condition, was significantly altered in the serum (Fig. S17).

Overall, these data demonstrate that *Bacteroides* in the gut leads to higher intestinal but not systemic propionate, the presence of *Bacteroides* also shifted the microbial community in the gut, particularly causing reduced relative abundance of *E. coli*, and the presence of *Bacteroides* in the gut has a greater impact on systemic (serum) pro-inflammatory cytokine levels than local (fecal) inflammatory marker levels, a result that has implications for the role of *Bacteroides* in the gut-lung axis in CF.

## DISCUSSION

Studies of pwCF have demonstrated that both *Bacteroides* and propionate are decreased in the stool (Antosca et al., 2019; Coffey et al., 2019; Vernocchi et al., 2018). Vernocchi *et al*. first identified decreased propionate in stool from children with CF. Similarly, Coffey *et al*. identified lower levels of pantetheine, a metabolite involved in propionate metabolism. Because *Bacteroides* is both a major component of a healthy microbiome and a large producer of propionate, it is likely that reduced *Bacteroides* is a factor in the observed reduction of intestinal propionate in pwCF, but this has not previously been demonstrated experimentally. Furthermore, pwCF are known to have higher levels of intestinal inflammation and propionate has been previously linked to decreases in the inflammatory response in non-CF intestinal epithelial cells (Bruzzese et al., 2014; Coffey et al., 2019; de Freitas et al., 2018; Iraporda et al., 2015; Parada Venegas et al., 2019).

The intestinal microbial composition has been linked to negative lung health outcomes in several CF clinical studies (Antosca et al., 2019; Bruzzese et al., 2014; Bruzzese et al., 2004; Burke et al., 2017; Coffey et al., 2019; de Freitas et al., 2018; Hayden et al., 2020; Hoen et al., 2015; Madan et al., 2012; Price & O’Toole, 2021). A recent study by Meeker *et al*. found that microbial colonization of germ-free CF mice (CFTR S489X) in part drives abnormal immune development (Meeker et al., 2020), demonstrating that dysbiosis in the CF intestine may be one driver of increased inflammation in pwCF.

We have shown here that *Bacteroides* secreted products downregulate the inflammatory response of CFTR-/- Caco-2 intestinal epithelial cells and that the production of propionate is required for this downregulation to occur. Deletion of the gene required for propionate production did not alter the production of any other major SCFAs, however, it is notable that this deletion causes *Bacteroides* to produce an increased amount of succinate, which in turn may affect production of other metabolites (Kovatcheva-Datchary et al., 2015). Succinate itself had no impact on IL-8 production *in vitro* (Fig. S10B). The finding that CFTR-/- Caco-2 cells are responsive to the anti-inflammatory effects of the three major SCFAs indicates that increasing intestinal SCFA concentrations, either through remediation of the gut microbiome or direct supplementation with SCFAs, has the potential to ameliorate intestinal inflammation for pwCF.

Interestingly, the *Bacteroides* isolates that we worked with here produced similar levels of propionate but showed differential capacities for IL-8 downregulation, indicating that propionate is not the sole contributor to IL-8 downregulation. *Bacteroides* isolates also produced low quantities of butyrate, a SCFA that to our knowledge has not previously been demonstrated to be produced by *Bacteroides*. Furthermore, the production of butyrate appears to be positively correlated with the anti-inflammatory effects of *Bacteroides* supernatants, further supporting our finding that butyrate production is biologically relevant and indicating that multiple metabolites may be responsible for the reduction in IL-8 levels we observe. We have identified two additional metabolites, lactate and riboflavin, which are detected in *Bacteroides* supernatants and have IL-8 down-regulatory effects. Thus, it is possible that a combination of these metabolites (and others) contributes to the anti-inflammatory effects of *Bacteroides*. Measurements of intestinal SCFAs have demonstrated that both propionate and butyrate are decreased in stool from pwCF (Coffey et al., 2019; Vernocchi et al., 2018). It is important to consider that *Bacteroides* are the primary source of some, but likely not all, of the important anti-inflammatory metabolites identified in this study. *Bacteroides* have been identified as the major intestinal producer of propionate, whereas members of the phylum Firmicutes are responsible for the bulk of intestinal butyrate production (Rios-Covian, Salazar, Gueimonde, & de Los Reyes-Gavilan, 2017). Interestingly, several butyrate-producing taxa, including *Faecalibacterium*, *Roseburia,* and *Bifidobacterium*, also have decreased relative abundance in the gut of pwCF (Price & O’Toole, 2021).

While we have focused on metabolites here, it is important to also consider the impacts of other *Bacteroides*-derived microbial products. We have shown that under our *in vitro* conditions, proteins do not apparently contribute to the anti-inflammatory effects of *Bacteroides* supernatants. However, it is possible that proteins not expressed under our laboratory conditions have important impacts on the immune system. Furthermore, we have not examined the impacts of polysaccharides in this study, which have previously been shown to be important in *Bacteroides*-host interactions. Perhaps the most well-known example is *Bacteroides* polysaccharide A (PSA), which is produced by *B. fragilis* and promotes proper T-cell subset development and promotes secretion of anti-inflammatory cytokine IL-10 by T-cells (Budden et al., 2017; Troy & Kasper, 2010).

Interestingly, capsule from *B. fragilis* has been shown to have anti-inflammatory effects in a mouse model of ulcerative colitis, an intestinal inflammatory disorder with similarities to CF intestinal manifestations (Zheng et al., 2020).

We have hypothesized a role for *Bacteroides* in local SCFA production, systemic inflammation, and/or lung health outcomes in CF. We therefore designed a mouse study to directly test these hypotheses whereby the native mouse microbiome in *Cftr^F508del^* mice was depleted through antibiotic treatment and then replaced by oral gavage with a pooled stool sample from children with CF. We have shown here that it is possible to replace the native mouse microbiome with a CF-like microbiome, and that when this microbiome is supplemented with *Bacteroides* isolates, these microbes will be maintained at high relative abundance for at least two weeks. As predicted from our *in vitro* studies, the presence of high *Bacteroides* caused an increase in measurable stool propionate. The mice with high *Bacteroides* also had significantly lower quantities of *systemic* levels of the pro-inflammatory cytokines KC (CXCL1), IL-5, IL-6, and MIP-1b.

In this model, we did not see direct effects on the airway. It is also important to note that while supplementation of mouse gut microbiota with *Bacteroides* results in an increased level of propionate in the stool of mice in our studies, the non-supplemented mice still produced detectable propionate despite low measured levels of *Bacteroides* by 16S rRNA gene sequencing. These data raise interesting implications that need to be more fully explored. First, while propionate drives reduction of IL-8 *in vitro*, as discussed above, we identified 3 other metabolites produced by *Bacteroides* that have a similar activity, and in fact, butyrate levels best correlate with the reduction in IL-8 levels *in vitro*. Thus, it may be the unusual suite of metabolites produced by *Bacteroides* spp. that together most effectively modulate systemic production of cytokines. It is also possible that *Bacteroides* spp. occupy a specific spatial niche that allows these microbes to most effectively communicate with the immune machinery. That is, the propionate we detected in the (-) *Bacteroides* condition in our analysis is not produced at the right time or in the right place by the other members of the microbiota to influence serum cytokine levels.

As mentioned above, we noted that while we can detect increased propionate in the stool in our mouse studies, we do not observe an increase of this SCFA in the serum. This is not unexpected, as under normal conditions propionate enters the portal vein and is metabolized in the liver such that only low concentrations of propionate are typically detectable in the periphery (Koh, De Vadder, Kovatcheva-Datchary, & Bäckhed, 2016). In contrast, in mice supplemented with *Bacteroides*, most of the observed impacts on cytokines were in the serum. An important implication of this finding is that it argues against a model whereby circulating SCFA are acting to impact the host. One simple model to reconcile these findings is that *Bacteroides* acts locally (perhaps via secreted metabolites) to modulate serum cytokine levels, and thus systemic inflammation.

Interestingly, the absence of *Bacteroides* was associated with a significant increase in the relative abundance of ASVs assigned to *Escherichia-Shigella*. An increased relative abundance of *E. coli* has also been detected in the gut microbiome of pwCF (Coffey et al., 2019; Hayden et al., 2020; Hoffman et al., 2014; Kristensen et al., 2020; Manor et al., 2016; Nielsen et al., 2016). Furthermore, previous work in a gnotobiotic mouse model has demonstrated that colonization with enterohemorrhagic *E. coli* is inhibited by pre-colonization with *B. frailis* or *B. vulgatus* (Saito et al., 2019). In a separate study, *E. coli* O157:H7 was shown to be inhibited by small organic acids (Shin, Suzuki, & Morishita, 2002). These results raise the intriguing possibility that the loss of *Bacteroides* and the propionate it produces might have implications for interspecies interactions in the context of the microbiota of infants and children with CF, specifically by allowing the expansion of *E. coli*. However, an important caveat to our dataset and others that rely on relative abundance data is that we cannot make any claims about absolute abundance. That is, the addition of *Bacteroides* may significantly decrease the detected relative abundance of *E. coli* without influencing absolute abundance.

We did not detect any differences in the levels of calprotectin, a frequently used marker of intestinal inflammation, in the stool from mice +/-*Bacteroides*. While studies have shown that gut microbiome composition is associated with lung health outcomes, evidence is mixed as to whether intestinal calprotectin is correlated with systemic outcomes (Price & O’Toole, 2021). The results reported here support the idea that while calprotectin may be a useful marker for local inflammation, it may not be as useful for assessing inflammation outside of the gut or for a tool to explore the gut-lung axis in mice.

This study highlights the importance of microbially-derived propionate and its role in maintaining intestinal homeostasis. It also demonstrates the potential for supplemented intestinal *Bacteroides* (i.e., as a probiotic) to modulate systemic inflammation. Finally, it shows that microbiome studies in *Cftr^F508del^* mice have the potential to better elucidate the underlying mechanisms in correlative, clinical microbiome studies.

## ACKNOWLEDGEMENTS

This work was supported by grants from the Cystic Fibrosis Foundation (OTOOLE19GO to G.A.O. and MADAN596389 to J.C.M.) and from the NIH (T32 AI007363 to C.E.P). Additional support was provided by the CF-Research Development Program (STANTO19R0), as well as DartCF and the BBC (P30-DK117469). We thank Mitch Drumm for generously providing the CRISPR-modified wild-type and CFTR-/- Caco-2 intestinal epithelial cells, and Craig Hodges for providing the *Cftr^F508del^* mouse line. We also thank Eric Martens for generously providing the *B. thetaiotaomicron* Δ*prp* and parent strains. We thank Joseph Sevigny for assistance with whole genome sequencing assembly.

## MATERIALS AND METHODS

### Cell Lines

CFTR was inactivated in a Caco-2 intestinal epithelial cell line by CRISPR/Cas9 targeting of Exon 11 (Hao et al., 2020); three such cell lines designated “S1”, “C9”, and “N5”, along with WT lines that were treated with a mock CRISPR/Cas9 protocol and designated “Y15”, “Y4”, and “N14”. These lines were generously provided by Mitch Drumm at Case Western Reserve University. The S1 cells are referred to as “CFTR-/CFTR- Caco-2 cells” in the text, and all experiments were performed in the S1 background unless otherwise noted. This cell line was maintained as previously described (Antosca et al., 2019). Briefly, cells were cultured at 37°C, 5% CO_2_ in Advanced MEM (Gibco #12492013) supplemented with 2 mM Glutamax (Gibco #35050061), penicillin-streptomycin (Gibco 50,000 U/500 ml #5140122), and 10% fetal bovine serum (FBS; Gibco; # 15140122). At passaging, cells were seeded into 24-well plates (Falcon #353047) at 100,000 cells/well or 96-well plates (Falcon #353072) at 10,000 cells/well. Cells in 96-well plates were grown for 1 week prior to use. Cells in 24-well plates were grown for 2 weeks prior to use.

### Bacterial Culture and Supernatant Preparation

*Bacteroides* isolates were regularly cultured in anaerobic boxes with GasPak EZ pouch system (BD Biosciences) on blood agar, which contained tryptic soy agar (BD Biosciences) + 5% sheep’s blood (Northeast Laboratory Services; Waterville NH). Bacterial supernatants were prepared by inoculating *Bacteroides* grown on blood agar into 5mL freshly prepared MEM (Gibco #51200038) supplemented with components from *Bacteroides* minimal medium (Martens, Chiang, & Gordon, 2008) and growing anaerobically for 24 hours. *Bacteroides* minimal medium supplements added to MEM are 4mM L-cysteine, 5ug/ml hemin chloride, 100μM MgCl_2_, 1.4μM FeSO4·7H2O, 50μM CaCl_2_, 1μg/ml vitamin K3 and 5ng/ml vitamin B12. This medium is referred to as supplemented MEM (sMEM) throughout the text. Oxygen was removed from the liquid medium by de-gassing in a Balch tube with 95% N2/5% CO_2_. Cultures were capped, crimped, and grown standing at 37°C for 24 hours. The cultures were collected and plated for colony forming units (CFUs) on blood agar. Supernatants were prepared by centrifuging the culture for 10 minutes at 4°C at 2823 x *g*, then sterile filtered with a 0.22uM filter to remove remaining bacteria. Supernatants were stored at -80°C prior to use in coculture assays or for quantification of metabolites.

### Coculture Assays

The CFTR-/CFTR-Caco-2 cells were washed 2X with MEM. *Bacteroides* supernatants were diluted 1:1 with fresh sMEM. As a control, freeze-thawed sMEM was also diluted 1:1 with fresh sMEM. Metabolite coculture assays were performed at the specified concentrations of each metabolite in MEM + L-gln. IL-1β was added to each condition at a final concentration of 10ng/mL to stimulate production of IL-8. 250uL or 100uL of sterile supernatants, or ½ freeze-thawed and ½ fresh sMEM as control, were added to CFTR-/CFTR-Caco-2 cells in 24 or 96-well plates, respectively. After incubation for 6 or 24 hours, as indicated in each experiment, supernatants were removed and centrifuged at 14,000 x *g* for 3 minutes at 4°C. Supernatant from the coculture experiment with CFTR-/CFTR- Caco-2 cells was transferred to a clean plastic 96-well plate and stored at -80°C; IL-8 was subsequently quantified by ELISA (Biolegend), as reported (Antosca et al., 2019).

### Viability Assay

To assess the viability of the CFTR-/CFTR- Caco-2 cells under the various treatment conditions assayed we used an XTT viability assay. XTT was prepared fresh prior to each experiment; 0.5mg/mL XTT was dissolved in PBS++ (Dulbecco’s PBS +1mM MgCl_2_ +0.1mM CaCl_2_, pH 8.2) for at least 30 minutes at 55°C. After cooling to room temperature, 0.025mM menadione was added. After supernatants were removed from the Caco-2 intestinal epithelial cells, the cells were washed 1X with PBS++, and then 250uL (24-well plates) or 100uL (96-well plates) XTT was added to each well. Plates were incubated at 37°C 5% CO_2_ for at least 3 hours or until orange color developed. 100uL was transferred from each well to a 96-well plate, OD450 was measured, and viability was calculated as a percentage of the IL-1β stimulated control.

### Fractionation and Proteinase K Treatment

Supernatants were prepared from *B. vulgatus* isolate CFPLTA003_2B as described above. Proteinase K (20mg/mL, VWR #E195) was added to the supernatant at 2.5uL/mL, the tube sealed, then the samples were incubated at 37°C for 1-2 hours, and then at 95°C for 15 minutes to inactivate proteinase K. A mock sample was incubated at the same temperatures for the same amount of time without Proteinase K. For fractionation experiments, 500uL supernatant was applied to 3 kDa size cut-off columns (Amicon Ultra #UFC5000324). Flowthrough and retentate fractions were collected according to the manufacturer’s instructions. The upper retentate was re-diluted to the original volume in fresh sMEM. After processing, standard coculture assays were performed as described above.

### Metabolite Quantification

#### In supernatants

Undiluted *Bacteroides* supernatants were stored frozen at -80°C prior to metabolite quantification. Frozen samples were shipped to Metabolon for analysis by ultra high-performance liquid chromatography/tandem accurate mass spectrometry (UHPLC/MS/MS) (Metabolon, Inc; Durham, NC). Data were re-scaled to a median of 1 and undetected metabolites were set to the minimum. All statistical analyses were performed on scaled data. SCFAs were quantified by gas chromatography-mass spectrometry (Michigan State University Mass Spectrometry and Metabolomics Core; protocol id MSU_MSMC_010). SCFA concentrations were calculated by normalization to standards.

#### In stool

Whole stool pellets were collected from each mouse, weighed, and immediately frozen at -80°C. Two metal beads and 600uL of methanol were added to each pellet and vortexed for 5 minutes. After a brief centrifugation step, 150uL of extract was analyzed by gas chromatography-mass spectrometry (Michigan State University Mass Spectrometry and Metabolomics Core; protocol id MSU_MSMC_010). Results were normalized to sample weight at collection.

#### In serum

12 SCFAs (C2-C8) were analyzed in serum by LC-MS/MS on a Waters Xevo TQ-S mass spectrometer at the Duke proteomics and metabolomics shared resource center. Data analysis was performed with Skyline software (www.skyline.ms). See supplemental methods for additional details.

### Genomic Analyses

Whole genome sequence information is available under bioproject accession PRJNA557692. Genomes were annotated and protein amino acid files generated with prokka (Seemann, 2014). Genus and species were determined by Sanger sequencing of the 16S amplicon for each strain (Forward primer: AGA GTT TGA TCC TGG CTC AG; Reverse primer: ACG GGC GGT GTG TRC). For strains with whole genome sequencing available, genus and species were confirmed by constructing an amino acid phylogeny including representative *Bacteroides* and *Parabacteroides* strains with known species assignments.

### *Bacteroides fragilis* toxin search

To determine whether any *B. fragilis* isolates encoded a gene for the *B. fragilis* enterotoxin (BFT), we first retrieved the Bft amino acid sequence from Uniprot (Q6UCA5). We then searched all completed *B. fragilis* genomes on the Integrated Microbial Genomes and Microbiomes system to identify homologs to the Uniprot Bft sequence (Chen et al., 2021). Five genes were identified (Gene IDs: 2649856759, 2649860343, 2649856758, 2848002094, 2649860342) across the 8 organisms searched, and amino acid sequences were downloaded for each gene. An align sequences protein BLAST was used to search for similarity of each amino acid sequence identified above against each *B. fragilis* genome with completed whole genome sequencing (PRJNA557692) in this study. Of the six *B. fragilis* isolates in this study, only one (AD126T_3B) did not have available sequencing data.

### Mouse Studies

C57BL/6 *Cftr^F508del^* mice deficient in production of the CFTR protein were obtained from Case Western Reserve (Loeven et al., 2021) and were treated with antibiotics for 21 days as previously described to suppress the endogenous intestinal microflora (Staley et al., 2017). After antibiotic treatment, mice were orally gavaged with 10uL stool pool from children with CF per gram of mouse weight. Stool pools contained either PBS or spiked-in *Bacteroides* from strains CFPLTA003_2B, SMC7758, and AD126T 3B. Each strain was grown anaerobically for 48 hours as a lawn on 10 blood agar plates and collected fresh for addition to stool pools. Collected *Bacteroides* were washed and re-suspended in PBS and an equal volume of PBS only was added to the (-) *Bacteroides* stool pool. The (+) *Bacteroides* condition contained a total inoculum of ∼10^9^-10^10^ CFU/mL *Bacteroides* and the (-) Bacteroides condition contained ∼10^4^ CFU/mL. Stool was regularly collected from mice before antibiotic treatment and for two weeks after oral gavage for analysis of microbiome composition (16S amplicon sequencing, UNH), cytokines (Luminex 32-plex; see Supplemental Materials 1), IgA (ELISA), and SCFAs (GC-MS, Michigan State University). Mice were then anesthetized with isoflurane and oropharyngeally challenged with 0.2mg/kg (5μg/25g) of LPS from *P. aeruginosa* 10 (Sigma). Mice were sacrificed with Euthasol 24-hours post-challenge. At sacrifice, whole lungs and intestinal tissue from the cecum, ileum, and colon were collected and immediately fixed in 10% buffered formalin. Whole blood was collected at the time of sacrifice. Serum was separated by allowing samples to stand at room temperature for 1-2 hours, and centrifuged at 1500g 4°C for 10 minutes. The serum (upper) fraction was removed and stored at -80°C prior to analysis.

### Ethics Statement

Mouse studies were performed in accordance with the Guide for the Care and Use of Laboratory Animals of the National Institutes of Health (NIH). The protocol was reviewed and approved (0002184) by the Institutional Animal Care and Use Committee (IACUC) at Dartmouth College. The Dartmouth College animal program is accredited with the Association for Assessment and Accreditation of Laboratory Animal Care International (AAALAC) under accreditation number D16-00166, and registered with the US Department of Agriculture (USDA), certificate number 12-R- 0001, and operates in accordance with Animal Welfare Assurance (NIH/PHS) under assurance number D16-00166 (A3259-01). A total of 26 mice (9 females and 17 males) 10-12 weeks of age at the first antibiotic treatment were included in the study, with 13 mice per condition (Supplemental Table 1). The experiment was independently performed 3 times. Animals were matched by age, sex, and body weight to the extent possible. Stool was collected from all mice, but serum was collected only for two of the three experiments (21 mice). Two mice in the (-) *Bacteroides* condition were excluded due to high relative abundance of *Bacteroides* in the stool, indicating possible cross contamination.

## Notes

### Competing Interest Statement

The authors have declared no competing interest.

