## Supplemental Methods for "Intestinal *Bacteroides* Modulates Systemic Inflammation and the Microbial Ecology in a Mouse Model of CF: Evidence for Propionate and other Short Chain Fatty Acids Reducing Systemic Inflammatory Cytokines"

**UPLC-MS/MS Analysis of Short Chain Fatty Acids in Mouse Serum Samples**

**DPMCF Project 5823**

***In collaboration with Courtney Price, Dartmouth College***

August 23, 2021

**Objective**: Measure short chain fatty acids (SCFAs) levels in serum samples from mice using LC-MS/MS with Multiple Reaction Monitoring.

**Duke Proteomics and Metabolomics Core Facility Contributors**: Laura Dubois (sample preparation, data collection, data analysis, report writing), Hannah Simmons (sample preparation, data review, report writing), and M. Arthur Moseley (scientific oversight).

**Short Summary:**

- 26 mouse serum samples were provided to the DPMSR and analyzed using a custom SCFA assay developed to quantitate 12 short chain fatty acids from acetic acid to octanoic acid (C2-C8). Calibrators were prepared from 0.098 µM-1 mM (0.98 µM-10 mM for C2).
    - Effective calibration range with dilution for serum is 0.029 µM-300 µM (0.29 µM-3 mM for C2).
- Data collection was performed using LC-MS/MS on a Waters Xevo TQ-S mass spectrometer, including calibration curves for each analyte in a manner consistent with the FDA Guidance for Bioanalytical Method Validation. These analyses are “For Research Purposes Only”.
- Data analysis was done using Skyline software (www.skyline.ms), and concentrations reported in µM for all samples.
- The final data is reported in the document “5823_results_081921.xlsx” as µM for serum in the tab **S3. Data Workup (uM)**.
- Blank sample (control) indicates some background from reagents in SCFA species, but this would be expected to be consistent across all samples and should not affect any statistical observations between groups. The concentrations in the blank were calculated in the same manner as in the study samples; thus these values can be subtracted from any samples.

### Introduction:

The SCFA method quantifies 12 SCFAs that can be present in biological samples (Table 1). The method is based on validated work published by Han et al. (2015) and Christoffersen (2017), and it was implemented in the Duke Proteomics and Metabolomics Core Facility (DPMCF) for the quantification of these 12 analytes in fecal and plasma/serum samples. The analytical protocol includes the requisite calibration standards and stable isotope labeled internal standards (SIS). The use of these standards facilitates harmonization and standardization within a project and across projects with the highest sensitivity possible. Selective analyte detection is accomplished using a triple quadrupole tandem mass spectrometer operated in Multiple Reaction Monitoring (MRM) mode, in which specific precursor to product ion transitions are measured for every analyte and their SIS. A representative UPLC-MS/MS chromatogram is presented in Figure 1 (A).

### Preparation of Stable-Isotope Internal Standard (SIS) Solution:

The SIS solution was prepared by reacting a standard mixture of the 12 SCFAs with $^{13}C_6$ labeled 3-nitrophenyl hydrazine(3-NPH, IsoSciences) at 200 mM and 120 mM N-(3-dimethylaminopropyl)-N'-ethylcarbodiimide hydrochloride (EDC) in 50/50 v/v EtOH/water. After 30 minutes at 35°C, the reaction was quenched by cooling the solution to 0°C and diluting the derivatization with 10% ethanol in water. Reaction of 3-NPH with the acids was catalyzed by EDC to form at hydrazone product, which is very stable.

A standard solution of the 12 acids was prepared with acetic acid at 1.0 mM and with all other acids at 0.10 mM. This solution was diluted in a ratio of 1 to 10 (v/v) with 50% ethanol. The $^{13}C_6$ labeled 3-nitrophenyl hydrazine hydrochloride standard is received as a 1 mg sample, $^{13}C_6$-3NPH (IsoSciences), which was combined with 50% ethanol. 120 mM EDC-6% pyridine solution was combined with the standard solution of the 12 acids and reacted for 30min at 35 °C and then quenched by cooling the solution to 0°C and diluting the derivative with 10% ethanol in water.

### Preparation of Calibration Standards:

A mixture of the C2-C8 short chain fatty acids was prepared volumetrically by dissolving the SCFAs in 50% ethanol in water for use as a stock solution to create a standard curve. The appropriate volumes were added so that acetic acid would be 1.0 mM and all other acids were 0.10 mM. The stock solution was then diluted serially to produce a 12-point calibration curve from 10 mM to 0.975 uM for acetic acid, and 1 mM to 0.098 uM for the C3-C8 short chain fatty acids. 20 uL of these standards was prepared in a 96-well plate alongside the samples using the same method.

### Sample Preparation:

Samples were prepared using the SOP for SCFA analysis developed in the DPMCF laboratory. First, the samples were transferred from plates to 0.5ml Eppendorf tubes. Sample duplicates from multiple wells were combined into one sample tube. 20ul of each sample was combined with 40uL of 100% ethanol and then the 26 sample extracts were centrifuged at 15,000 rpm for 15 minutes at 4C. An equal volume from all samples was pipetted out and combined to make a study pool QC (SPQC). 20 uL from each sample was derivatized by adding 20 uL of 200 mM 3-nitrophenyl hydrazine (3-NPH) in 50% ethanol with 6% pyridine and 120 mM N-(3-dimethylaminopropyl)-N'-ethylcarbodiimide hydrochloride (EDC·HCl) in 50% ethanol, incubating at 35°C for 30 minutes while shaking. The reaction was quenched with 760 uL of cold 10% ethanol in water with 1% formic acid and held at 4 °C for the double blank, standards and QCs. The same quenching protocol was used for the SPQC, study samples and "serum-prep" blank except 40 uL of cold 10% ethanol/1% formic was used. The plate was spun at 3000 rpm for 5 minutes. A 25 ul aliquot of the reaction solution from each sample was transferred and combined with 25 uL of the stable isotope standards (SIS) solution. (The 100x concentrated SIS solution is diluted 100x in 10% ethanol/1% formic acid prior to addition). An aliquot of Golden West serum (3x diluted in 100% ethanol) was also extracted to be used for quality control. The plate was capped with a polypropylene cap mat and placed in the LC autosampler for immediate analysis.

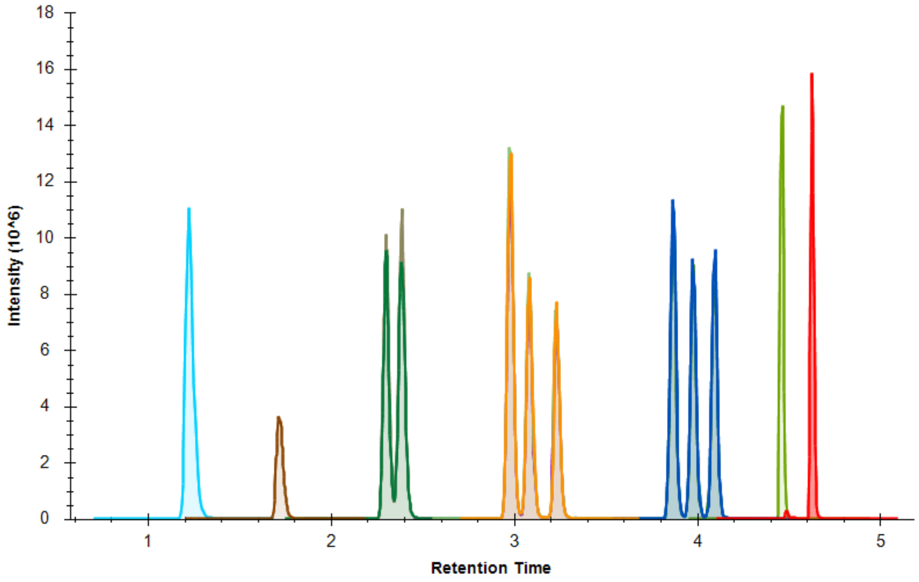


**Figure 1.** Short Chain Fatty Acid (SCFA) analysis by LC-MS/MS. The figure shows the overlaid chromatogram traces for each of the SCFA species, from Acetate (C2) to Octanoate (C8). This representative figure is from an equimolar mixture of standards. Each color represents the measurement of a SCFA with a specific number of carbons, and the separate peaks of each color are geometric isomers (where relevant).

### Sample Analysis by LC-MS/MS:

*Short Chain Fatty Acids.* UPLC separation of the SCFAs was performed using a Waters Acquity LC with a Waters Acquity 2.1 mm x 50 mm 1.7 µM BEH C18 column fitted with a Waters Acquity C18 1.7 µM Vanguard guard column. Analytes were separated using a gradient from 20% solvent B to 40% solvent B in 3.5 minutes followed by another ramp up to 100% solvent B over the next minute and then a hold. Solvents A & B were 0.1% formic acid in water and acetonitrile, respectively. The total UPLC analysis time was approximately 8 minutes. The method uses electrospray ionization in negative mode introduced into a Xevo TQ-S mass spectrometer (Waters) operating in the Multiple Reaction Monitoring (MRM) mode. MRM transitions (compound-specific precursor to product ion transitions) for each analyte and internal standard were collected over the appropriate retention time window. All the compounds measured in the MRM assay, including their retention times, precursor and product ions, and calibration ranges are included in **Table 1**. The calibration ranges listed are the pre-dilution factor values. The calibration ranges for serum post dilution factor are in S3 in the Excel data document. The separation of the individual SCFAs and the geometric isomers by RPLC is shown in **Figure 1**.

The standard curve and calibration QCs were run once at the beginning of the run queue and once at the end of the sample run queue. The study pool QCs (SPQC) and Golden West serum were run once at the beginning, once in the middle and once at the end of the run queue. The injection volume for each injection was 5 µL.

**Data Analysis**

All data was analyzed in Skyline v21.1.1 (www.skyline.ms) which includes raw data import, peak integration, and a linear regression fit with $1/x^2$ weighting for the calibration curves. The chromatography utilized shows very robust retention time reproducibility. The variation in retention time for the vast majority of the samples is +/- 0.01 minutes.

The calibration curves for all the analytes contain 12 points, from 0.98 uM to 10 mM for AA, and from 0.098 to 1 mM for the other analytes. Each calibrator's residual bias was calculated after regression fit, and any point other than the lower and upper limits of quantitation where the residual fell outside of 15% were removed from the calibration curve equation. The lower limit of quantitation (LLOQ) was defined to be the lowest point that meets the criteria of having a bias < 20% and the upper limit of quantitation (ULOQ) is defined in the same way for the highest point. The calibration range that was used for concentration calculations in the samples is listed in Table 1 for each analyte. **Figure 2** provides an example calibration curve for valeric acid, characteristic of most analytes in the panel.

The Skyline file containing the data from the 12 analytes, with all samples and calibration curves, has been included in the data return as a zip archive "5823_results_081921.xlsx"

| Molecule Name | Abbreviation | Label Type | Precursor m/z | Product m/z | Retention Time | Calibration Range (uM) | Cal Curve R^2 |
|---|---|---|---|---|---|---|---|
| Acetic acid | AA | light | 194 | 152, 137 | 1.24+/-0.01 | 15.6-10000 | 0.9981 |
| | | heavy | 200 | 143 | 1.24+/-0.01 | | |
| Propionic acid | PA | light | 208 | 165, 137 | 1.73+/-0.01 | 0.0975-1000 | 0.9956 |
| | | heavy | 214 | 143 | 1.73+/-0.01 | | |
| iso-butyric acid | i-BA | light | 222 | 179, 137 | 2.3+/-0.01 | 0.195-1000 | 0.9947 |
| | | heavy | 228 | 143 | 2.3+/-0.01 | | |
| butyric acid | BA | light | 222 | 152, 137 | 2.39+/-0.01 | 0.78-1000 | 0.9980 |
| | | heavy | 228 | 143 | 2.39+/-0.01 | | |
| 2-methyl butyric acid | 2-Me-BA | light | 236 | 152, 137, 81 | 2.98+/-0.01 | 0.195-1000 | 0.9990 |
| | | heavy | 242 | 143 | 3+/-0.02, 2.97 +/-0.03 | | |
| iso-valeric acid | i-VA | light | 236 | 152, 137 | 3.08+/-0.01, 3.07 +/-0.02 | 0.0975-1000 | 0.9999 |
| | | heavy | 242 | 143 | 3.08+/-0.01 | | |
| valeric acid | VA | light | 236 | 152, 137 | 3.23 +/-0.01 | 0.0975-1000 | 1.0000 |
| | | heavy | 242 | 143 | 3.23 +/-0.01 | | |
| 3-methyl valeric acid | 3-Me-VA | light | 250 | 152, 137 | 3.86 +/-0.01, 3.84 +/-0.02 | 0.0975-1000 | 0.9952 |
| | | heavy | 256 | 143 | 3.86 +/-0.01 | | |
| iso-caproic acid | i-CA | light | 250 | 152, 137 | 3.97+/-0.01 | 0.0975-1000 | 0.9849 |
| | | heavy | 256 | 143 | 3.97+/-0.01 | | |
| caproic acid | CA | light | 250 | 152, 137 | 4.09+/-0.01 | 0.195-1000 | 0.9948 |
| | | heavy | 256 | 143 | 4.08+/-0.01 | | |
| Heptanoic acid | HA | light | 264 | 137 | 4.46+/-0.01 | 0.0975-200 | 0.9987 |
| | | heavy | 270 | 143 | 4.46+/-0.01 | | |
| Octanoic acid | OA | light | 278 | 137 | 4.63+/-0.01 | 0.195-200 | 0.9985 |
| | | heavy | 284 | 143 | 4.63+/-0.01 | | |

**Table 1.** Short chain fatty acid (SCFA) metabolites monitored using this method, along with their analytical characteristics (precursor m/z, product m/z, and retention time). Note that the m/z values include the 3-NPH derivatization. The unlabeled compounds are referred to as “light”, the $^{13}C_6$ labeled compounds from the SIS solution are “heavy”. The calibration range and $R^2$ value for the standard curves are also listed.

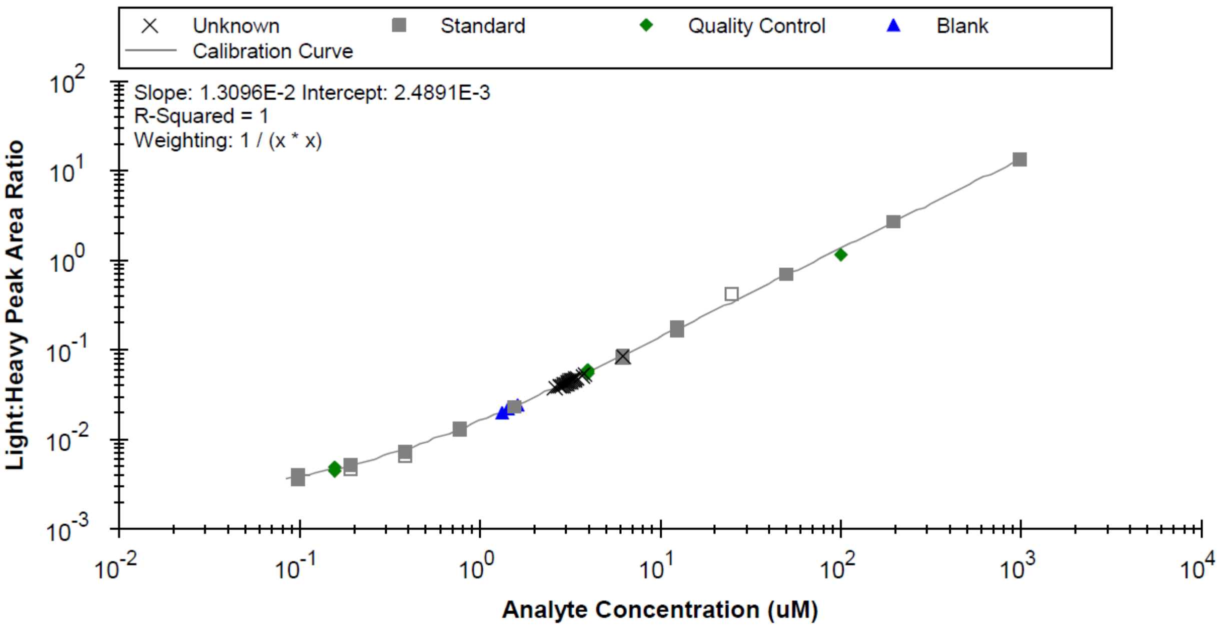


**Figure 2.** Example calibration curve (for C5, valeric acid) with study samples (marked with X's) spanning the middle of thedynamic range.

The table below (Table 2) shows the %CV for each analyte from the 3 injections of the SPQC.

**Reproducibility Metrics**

__26__ out of the 26 samples were included when creating the SPQC to run 3x across the run queue. The table below (Table 2) shows the percent coefficient of variation (CV%) for each analyte from the 3 injections of the SPQC.

| SPQC SCFA | Average (uM) | %CV |
|---|---|---|
| acetic acid | 545.923 | 0.40% |
| propionic acid | 9.206 | 1.77% |
| iso-butyric acid | 1.276 | 2.72% |
| butyric acid | 1.868 | 0.93% |
| 2-methyl butyric acid | 0.463 | 1.45% |
| iso-valeric acid | 0.497 | 1.01% |
| valeric acid | 0.966 | 3.76% |
| 3-methyl valeric acid | 0.087 | 4.04% |
| iso-caproic acid | 0.157 | 5.18% |
| caproic acid | 1.370 | 1.57% |
| heptanoic acid | 1.369 | 0.58% |
| octanoic acid | 2.044 | 0.88% |

**Table 2**. Average concentration of SCFA in the SPQC sample across 3 injections and the corresponding %CV with all less than 10%.

### Data Return Document Descriptions:

The Excel workbook uploaded to Express “5823_results_08192021” containing 4 worksheets.  A brief summary of each worksheet is included below:

**S1. Calibration Ranges:** same as Table 1 above

**S2. Raw Data:** All raw data from Skyline

**S3. Data Workup uM**: final data workup with serum SCFA concentrations in uM

**S4**. **SPQC Metrics:** same as Table 2 above
