## Supplemental Figures for "Intestinal *Bacteroides* Modulates Systemic Inflammation and the Microbial Ecology in a Mouse Model of CF: Evidence for Propionate and other Short Chain Fatty Acids Reducing Systemic Inflammatory Cytokines"

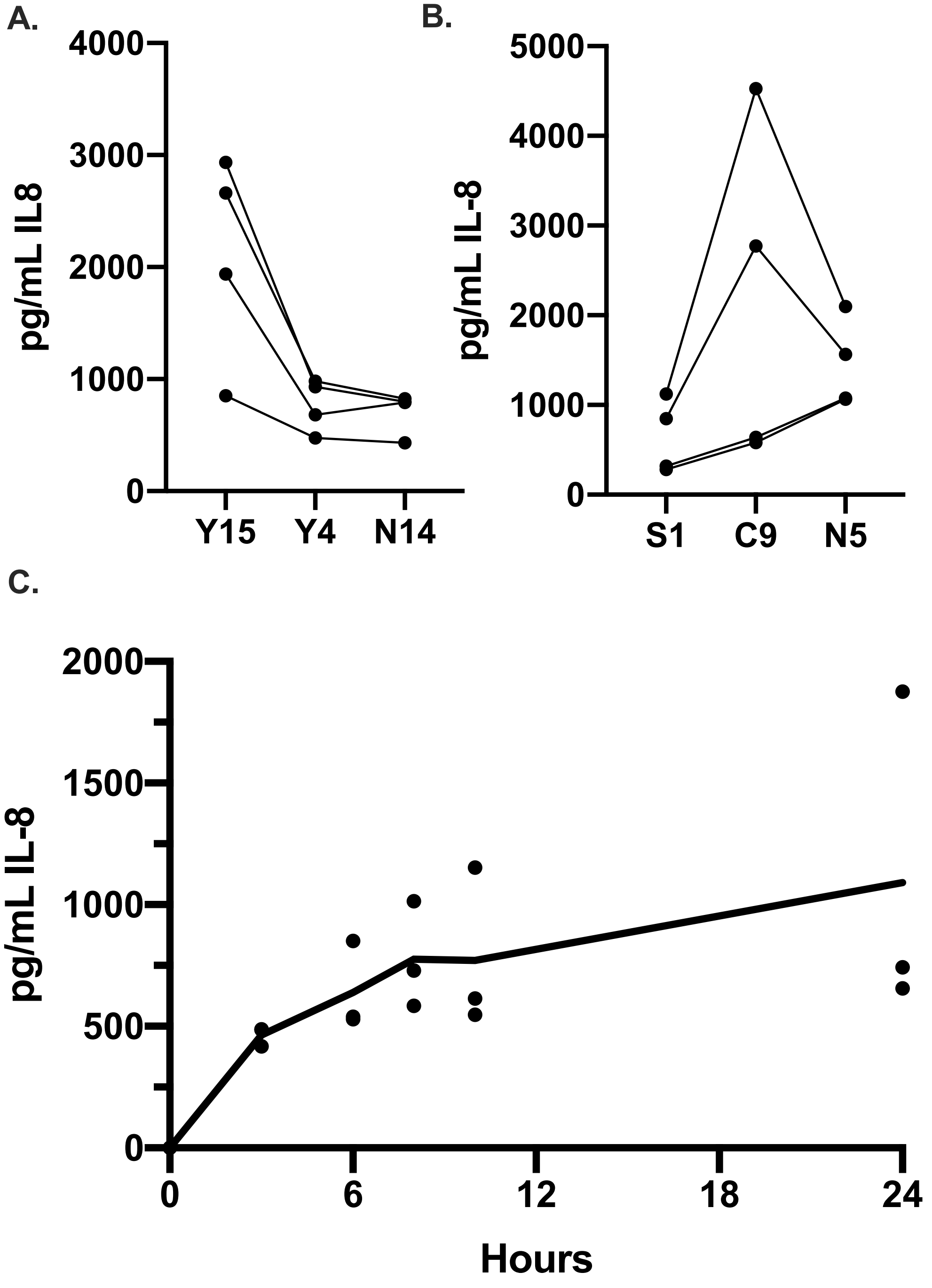


**Figure S1. IL-8 production by Caco-2 intestinal epithelial cells stimulated with IL-1β.** CFTR-/- Caco-2 intestinal epithelial cell lines were generated by CRISPR/Cas9 editing by Hao et al (Hao et al., 2020). Wild-type cell lines were generated from the same Caco-2 parent as the CFTR-/- lines, but these control lines were subjected to mock CRISPR/Cas9 treatment. The indicated Caco-2 cell lines were cultured for 2 weeks on plastic in 24-well plates and then treated with 10ng/mL IL-1β in MEM + L-gln medium. IL-8 was quantified by ELISA after 24 hours of IL-1β stimulation for three separate A) wild-type or B) CFTR-/- cell lines. Each point represents the average of four technical replicates from a single biological replicate (n=4), and 3-4 independent biological replicates were performed. Lines connect biological replicates in different cell lines that were performed on the same day. C) IL-8 was quantified by ELISA after 3, 6, 8, 10, and 24 hours of IL-1β exposure to the CFTR-/- S1 Caco-2 cells. Individual points indicate the average of three technical replicates from a single biological replicate (n =3 biological replicates), and the solid line indicates the average of three biological replicates at each time point.

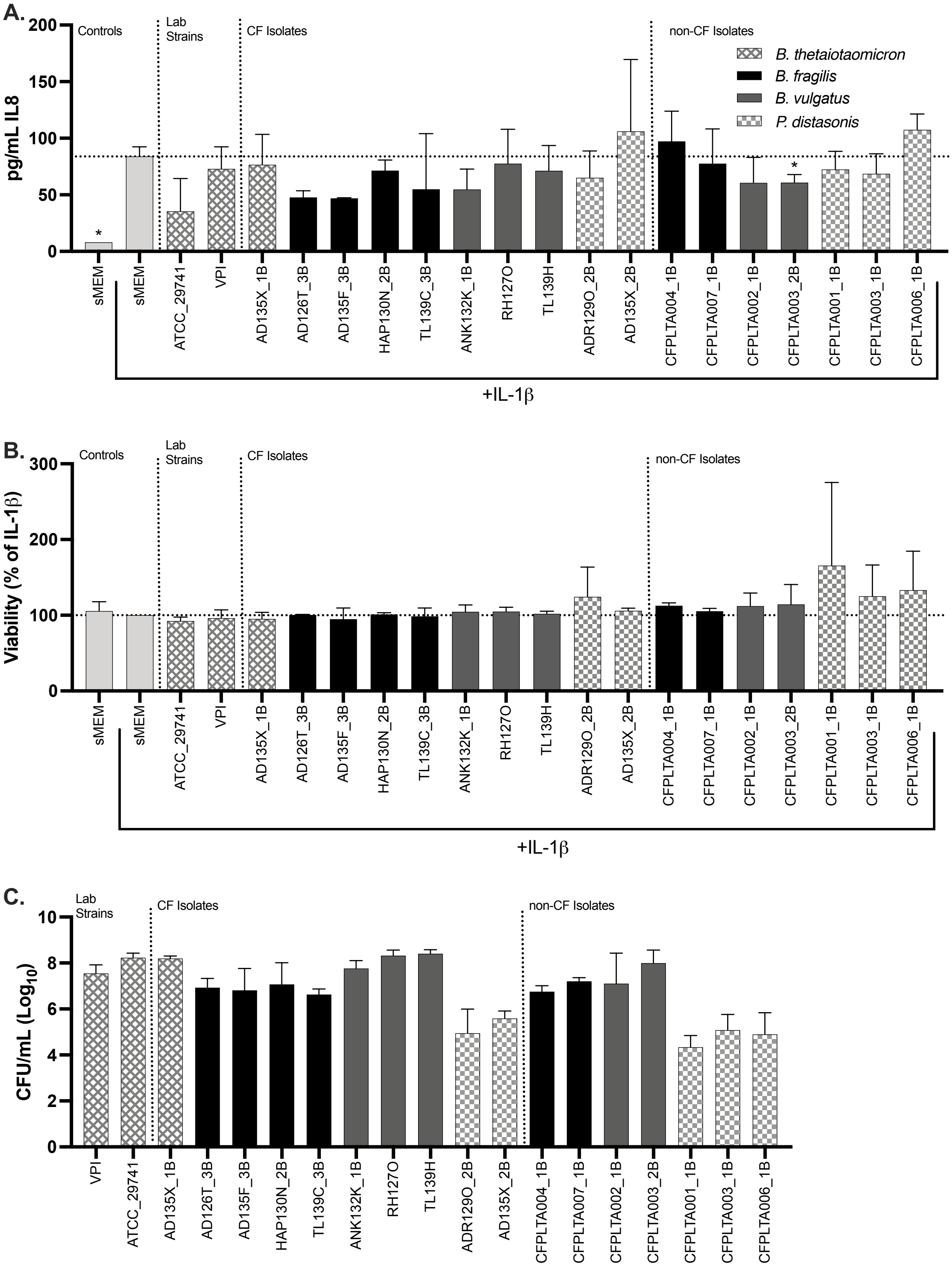

A.
B.
C.
Controls
Lab Strains
CF Isolates
non-CF Isolates
B. thetaiotaomicron
B. fragilis
B. vulgatus
P. distasonis
pg/mL IL8
Viability (% of IL-1β)
CFU/mL (Log10)
sMEM
ATCC_29741
VPI
AD135X_1B
AD126T_3B
AD135F_3B
HAP130N_2B
TL139C_3B
ANK132K_1B
RH127O
TL139H
ADR129O_2B
AD135X_2B
CFPLTA004_1B
CFPLTA007_1B
CFPLTA002_1B
CFPLTA003_2B
CFPLTA001_1B
CFPLTA003_1B
CFPLTA006_1B
+IL-1β

**Figure S2. *Bacteroides* isolate screen for effects on IL-8.** Cell-free, filter sterilized supernatants of the indicated *Bacteroides* strains were prepared in sMEM, applied to 1-week old CFTR-/- Caco-2 cells grown in 96 well plates, and then incubated for 6 hours as described in the Materials and Methods. Three biological replicates were performed for each isolate. The key in A applies to all graphs in the figure. A) IL-8 production was quantified by ELISA for CFTR-/- Caco-2 cells exposed to cell-free, filter sterilized supernatant from each isolate. Significance was tested by paired one-way ANOVA followed by Dunnett's post-test with MEM + IL-1$\beta$ as the reference. * $p < 0.05$. B) The XTT assay was used to quantify CFTR-/- Caco-2 cellular viability after exposure to the cell-free, filter sterilized supernatant from the indicated isolate. Significance was tested by paired one-way ANOVA followed by Dunnett's post-test with MEM + IL-1$\beta$ as the reference. * $p < 0.05$. C) Bacterial cultures were plated prior to removal of the bacterial cells and filter sterilization to quantify CFU/mL for each isolate.

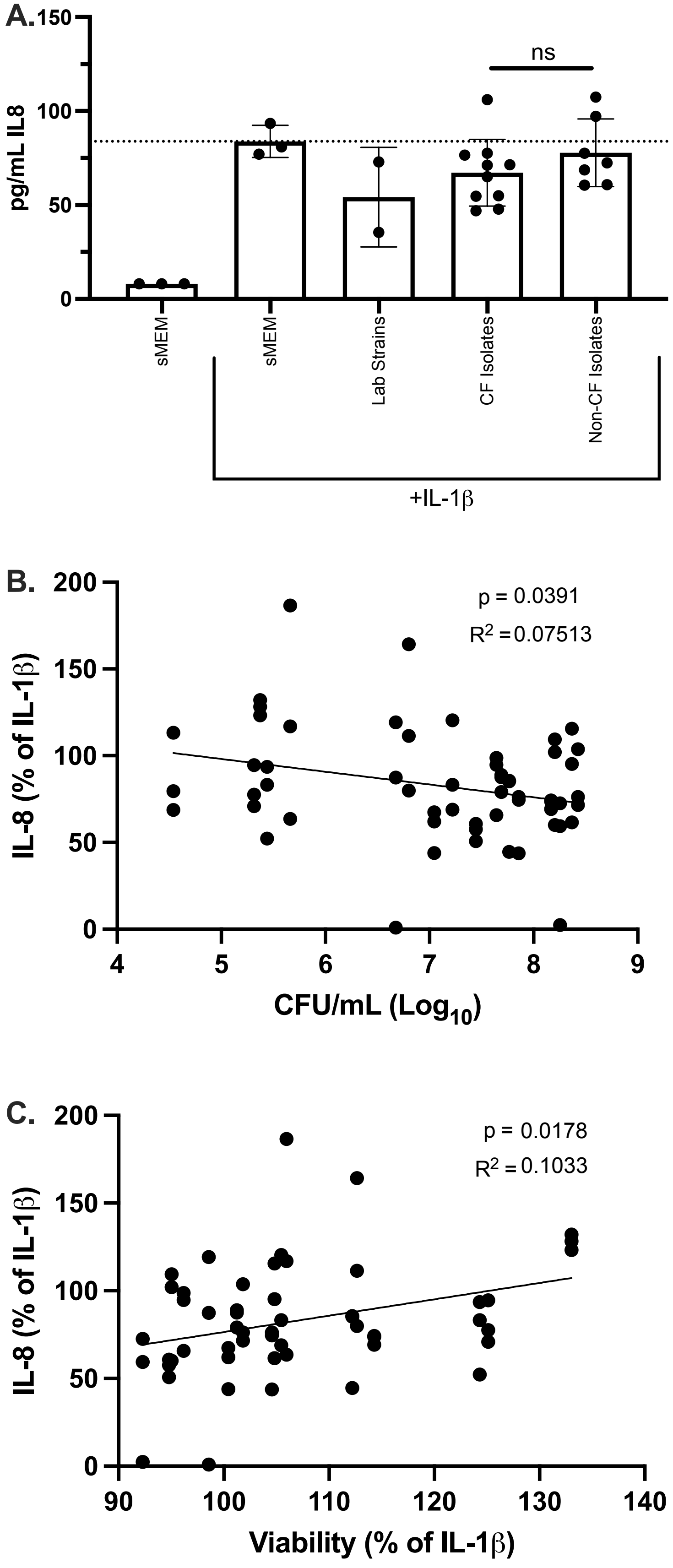
A.
150
100
50
0
pg/mL IL8
ns
sMEM
sMEM
Lab Strains
CF Isolates
Non-CF Isolates
+IL-1β
B.
200
150
100
50
0
4
5
6
7
8
9
p = 0.0391
$R^2$ = 0.07513
IL-8 (% of IL-1β)
CFU/mL ($Log_{10}$)
C.
200
150
100
50
0
90
100
110
120
130
140
p = 0.0178
$R^2$ = 0.1033
IL-8 (% of IL-1β)
Viability (% of IL-1β)

**Figure S3. *Bacteroides* isolate correlations with isolate origin, CFU/mL, and cellular viability.** Data in this graph are from the *Bacteroides* isolate screen used to generate the plots in Figure S2. A) Isolates were grouped by origin, and each point represent three biological replicates for a single isolate. An unpaired student's t-test was used to test for significant differences between the CF and non-CF isolates. B) Average $Log_{10}$ CFU/mL of each isolate was plotted versus IL-8 as a percentage of MEM + IL-1β control for each biological replicate. C) Average viability of the cell line in the presence of each isolate was plotted versus IL-8 as a percentage of MEM + IL-1β control for each biological replicate. B-C) Simple linear regression was used to calculate $R^2$ and p value.

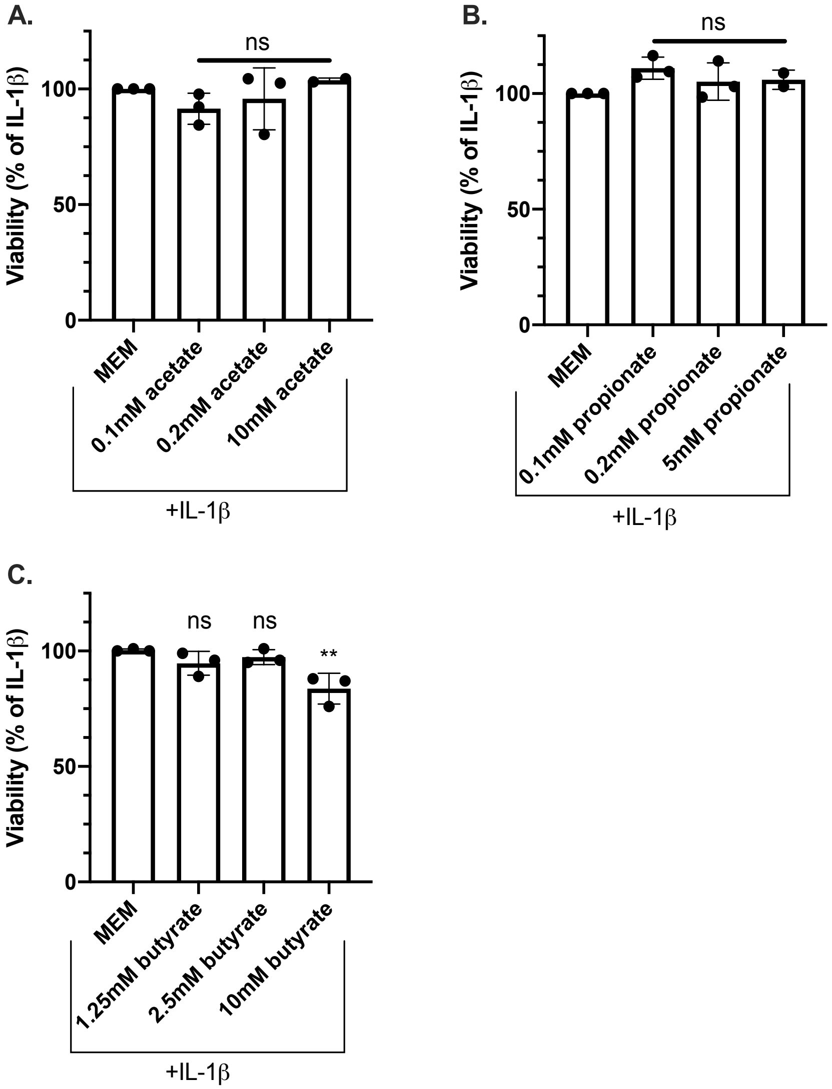


**Figure S4. Viability of CFTR-/- Caco-2 cell cultured with SCFAs.** CFTR-/- Caco-2 cells were cultured for 2 weeks in 24-well plates, and then exposed to MEM with the indicated SCFAs for 24 hrs. Viability was measured by XTT assay after exposure to A) sodium acetate B) sodium propionate and C) sodium butyrate at the indicated concentrations. The value displayed is a percentage of the viability of Caco-2 cells treated with MEM + IL-1β. Significance was tested by unpaired one-way ANOVA with Dunnett's post-test and MEM + IL-1β as the reference. **$p<0.01$.

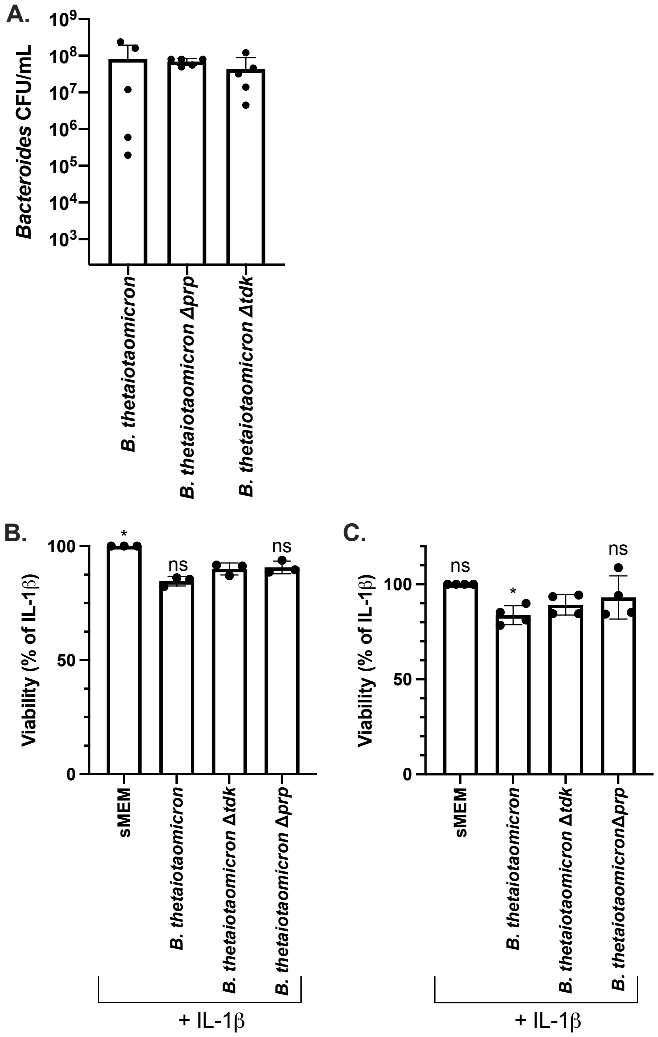


**Figure S5. *B. thetaiotaomicron* CFU/mL and impact on Caco-2 cell viability.** A) CFU/mL of cultures prepared for the indicates isolates. Each point represents a single biological replicate, with 5 total biological replicates performed. No significant differences were identified by one-way ANOVA followed by Tukey's post-test. Viability of the Caco-2 cells was measured by XTT assay after exposure to *B. thetaiotaomicron* supernatants for B) 6 hours and C) 24 hours. Significance was tested by unpaired one-way ANOVA followed by Dunnett's post-test with *B. thetaiotaomicron Δtdk* as the reference. *$p<0.05$.

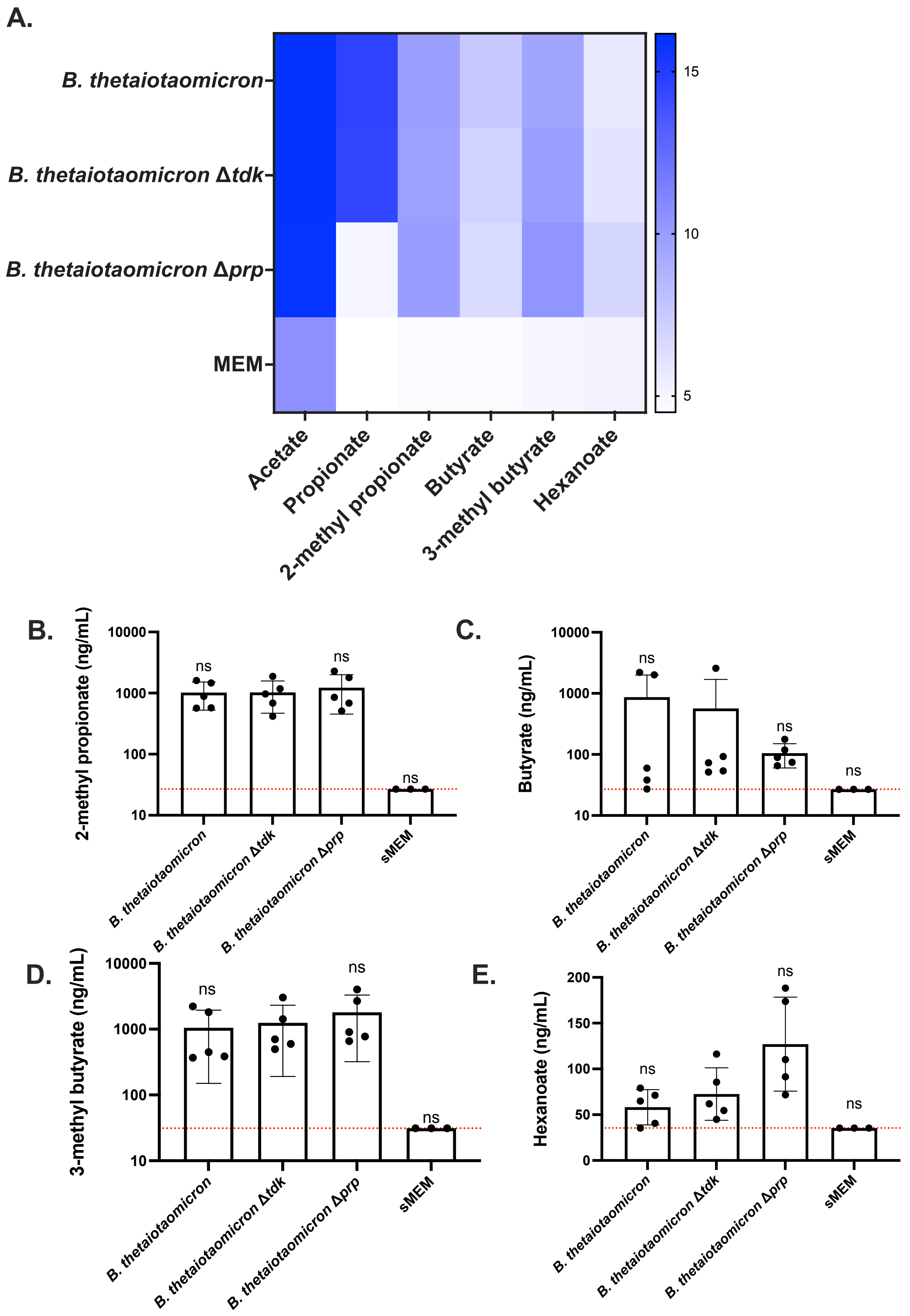

A.
B. thetaiotaomicron
B. thetaiotaomicron Δtdk
B. thetaiotaomicron Δprp
MEM
Acetate
Propionate
2-methyl propionate
Butyrate
3-methyl butyrate
Hexanoate
15
10
5
B.
2-methyl propionate (ng/mL)
C.
Butyrate (ng/mL)
D.
3-methyl butyrate (ng/mL)
E.
Hexanoate (ng/mL)
ns
sMEM
10000
1000
100
10
200
150
50
0

**Figure S6. Deletion of a propionate synthesis gene in *B. thetaiotaomicron* does not have detectable off-target effects.** SCFAs were quantified in undiluted, sterile-filtered supernatants by GC-MS. A) Heatmap of $\log_2$ transformed concentration (ng/mL) of each detectable SCFA. Pentanoic, 2-methyl pentanoic, and heptanoates were not detected above the LOD. B) 2-methyl propionate, C) butyrate, D) 3-methyl butyrate, and E) hexanoate were detected above background but were not significantly different between strains. Values below the limit of detection were thresholded to the LOD. Dashed red line indicates the LOD. Significance was tested by unpaired one-way ANOVA followed by Dunnett's post-test with *B. thetaiotaomicron* Δ*tdk* as the reference; ns = not significant.

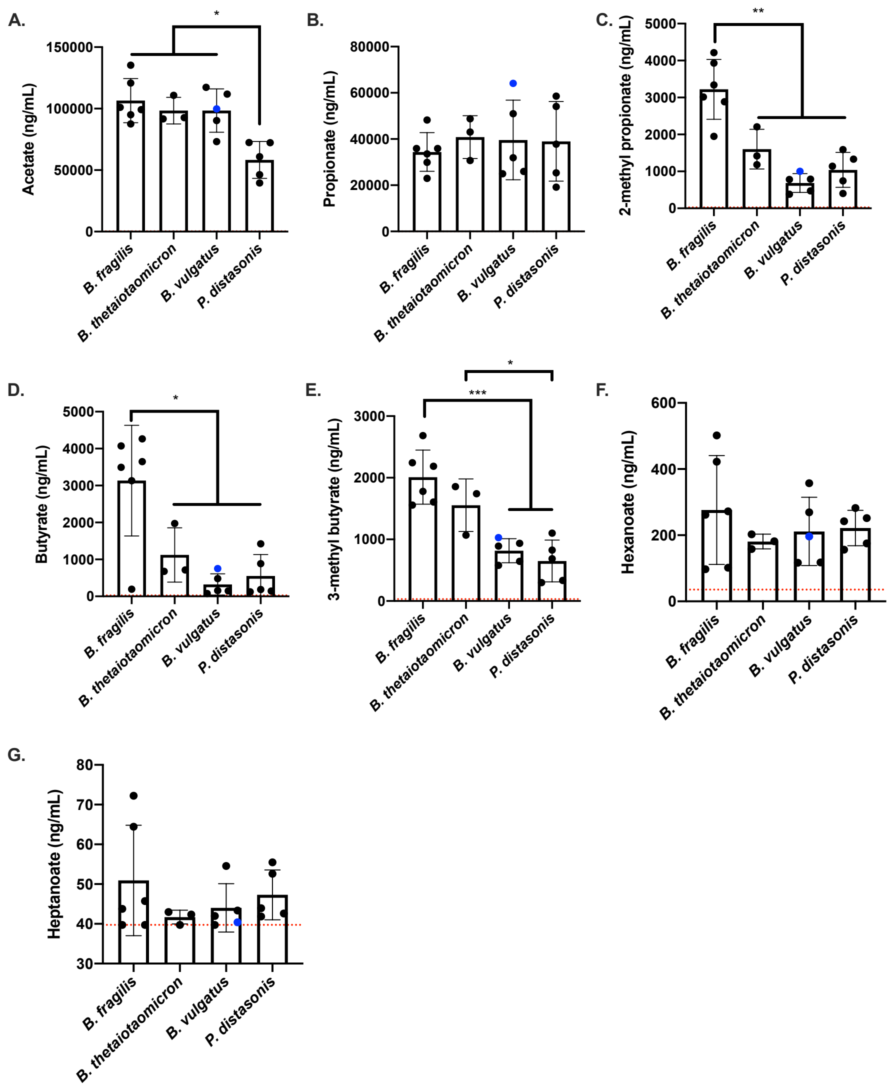


**Figure S7. SCFA quantification from *Bacteroides* isolate supernatants.** SCFAs were quantified in undiluted, sterile-filtered supernatants by GC-MS. Each point indicates the average SCFA concentration in ng/mL from 4-5 biological replicates from an individual isolate. The dashed red line indicates the LOD. Isolates are grouped in each graph by species. CFPLTA003_2B is colored blue in each graph for ease of comparison. Significance was tested by unpaired one-way ANOVA followed by Tukey's post-test. All significant pairwise differences are annotated. *p<0.05, **p<0.01, ***p<0.001.

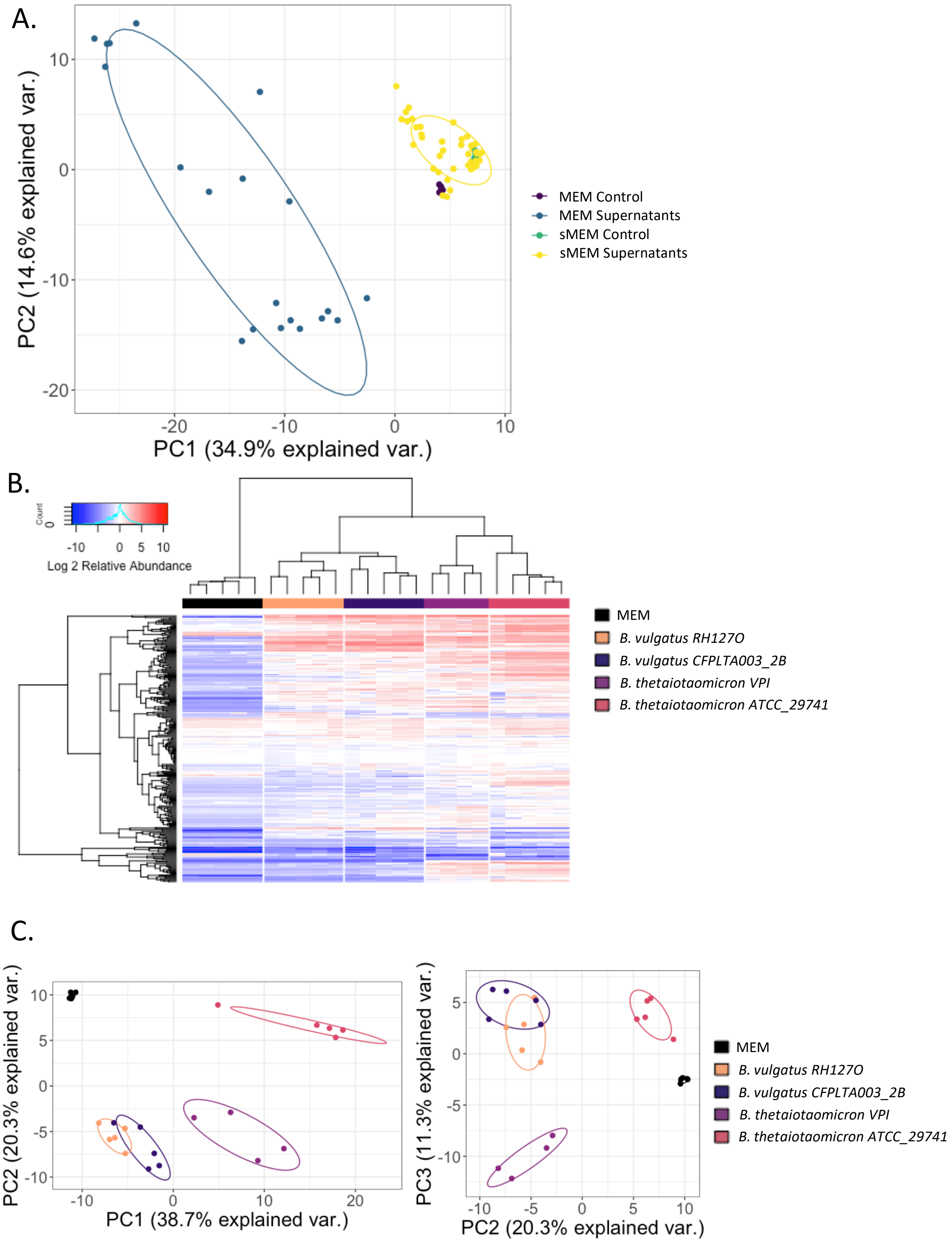
A.
PC1 (34.9% explained var.)
PC2 (14.6% explained var.)
MEM Control
MEM Supernatants
sMEM Control
sMEM Supernatants
B.
Count
Log 2 Relative Abundance
MEM
B. vulgatus RH127O
B. vulgatus CFPLTA003_2B
B. thetaiotaomicron VPI
B. thetaiotaomicron ATCC_29741
C.
PC1 (38.7% explained var.)
PC2 (20.3% explained var.)
PC2 (20.3% explained var.)
PC3 (11.3% explained var.)
MEM
B. vulgatus RH127O
B. vulgatus CFPLTA003_2B
B. thetaiotaomicron VPI
B. thetaiotaomicron ATCC_29741

**Figure S8. LC-MS quantification of *Bacteroides* and *Parabacteroides* supernatants.** *Bacteroides* isolates were cultured in MEM + L-gln for 48 hours or sMEM for 24 hours prior to metabolite quantification by LC-MS. A) PCA plot of all metabolites detected in both MEM and sMEM *Bacteroides* supernatants. Each point represents a single biological replicate and is color-coded by condition. B) Heatmap of all metabolites detected in supernatants from *Bacteroides* cultured in MEM. Each column represents a single biological replicate and is color-coded by strain. The heatmap was generated in R with the Heatmap.2 function. C) PCA plots of all metabolites detected in MEM *Bacteroides* supernatants. Left: PC1 vs PC2. Right: PC2 vs. PC3. Each point represents a single biological replicate and is color-coded by strain.

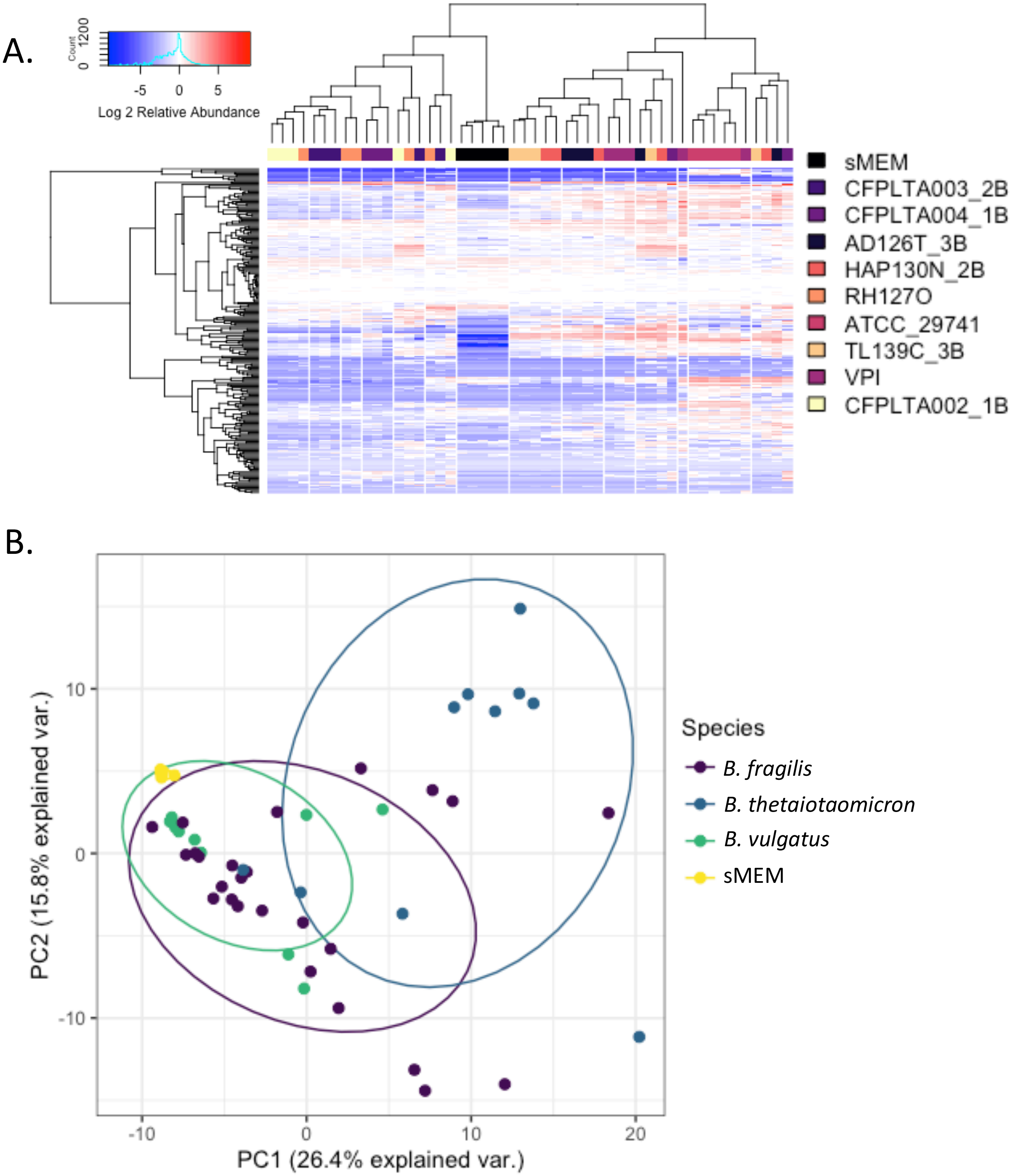
A.
Count
0
1200
-5
0
5
Log 2 Relative Abundance
sMEM
CFPLTA003_2B
CFPLTA004_1B
AD126T_3B
HAP130N_2B
RH127O
ATCC_29741
TL139C_3B
VPI
CFPLTA002_1B
B.
PC2 (15.8% explained var.)
PC1 (26.4% explained var.)
10
0
-10
-10
0
10
20
Species
B. fragilis
B. thetaiotaomicron
B. vulgatus
sMEM

**Figure S9. LC-MS quantification of *Bacteroides* and *Parabacteroides* supernatants cultured in sMEM.** *Bacteroides* isolates were cultured in sMEM for 24 hours prior to metabolite quantification by LC-MS. A) Heatmap of all metabolites detected in supernatants from *Bacteroides* cultured in sMEM. Each column represents a single biological replicate and is color-coded by strain. The heatmap was generated in R with the Heatmap.2 function. B) PCA plots of all metabolites detected in sMEM *Bacteroides* supernatants. Each point represents a single biological replicate and is color-coded by *Bacteroides* species.

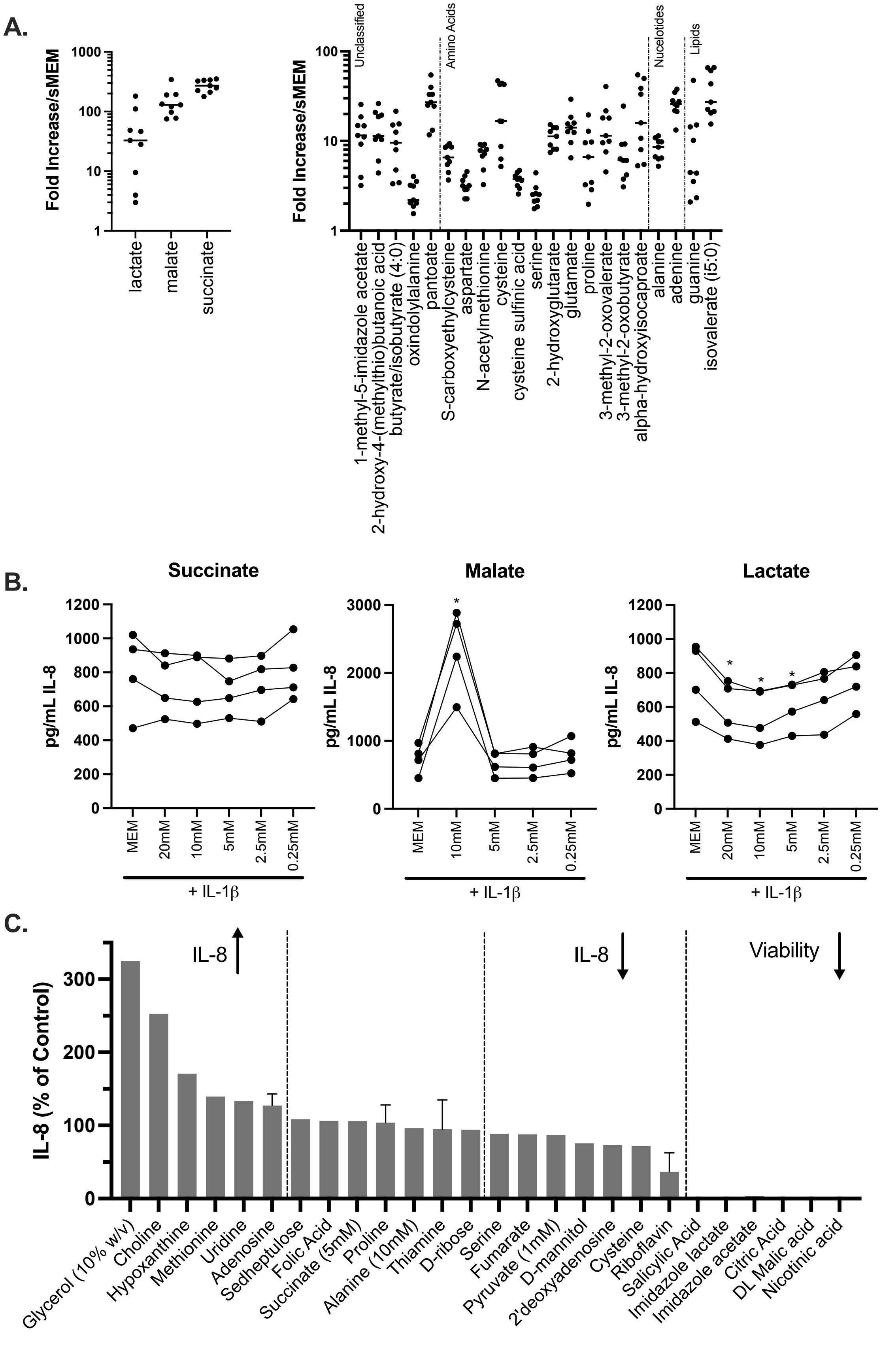
A.
Fold Increase/sMEM
1000
100
10
1
lactate
malate
succinate
Unclassified
Amino Acids
Nucelotides
Lipids
1-methyl-5-imidazole acetate
2-hydroxy-4-(methylthio)butanoic acid
butyrate/isobutyrate (4:0)
oxindolylalanine
pantoate
S-carboxyethylcysteine
aspartate
N-acetylmethionine
cysteine
cysteine sulfinic acid
serine
2-hydroxyglutarate
glutamate
proline
3-methyl-2-oxovalerate
3-methyl-2-oxobutyrate
alpha-hydroxyisocaproate
alanine
adenine
guanine
isovalerate (i5:0)
B.
Succinate
Malate
Lactate
pg/mL IL-8
MEM
20mM
10mM
5mM
2.5mM
0.25mM
+ IL-1β
C.
IL-8 (% of Control)
IL-8 ↑
IL-8 ↓
Viability ↓
Glycerol (10% w/v)
Choline
Hypoxanthine
Methionine
Uridine
Adenosine
Sedheptulose
Folic Acid
Succinate (5mM)
Proline
Alanine (10mM)
Thiamine
D-ribose
Serine
Fumarate
Pyruvate (1mM)
D-mannitol
2'deoxyadenosine
Cysteine
Riboflavin
Salicylic Acid
Imidazole lactate
Imidazole acetate
Citric Acid
DL Malic acid
Nicotinic acid

**Figure S10. Impacts of highly produced metabolites on IL-8.** LC-MS quantification of *Bacteroides* and *Parabacteroides* supernatants cultured in sMEM. *Bacteroides* isolates were cultured in sMEM for 24 hours prior to metabolite quantification by LC-MS. A) All metabolites detected >3-fold over sMEM control for every isolate and displayed as a ratio of the metabolite detected in supernatant/sMEM control. The panel on the left highlights SCFAs. B) CFTR-/- Caco-2 cells were cultured for 2 weeks in 24-well plates and then treated with IL-1β alone, or with the addition of the indicated metabolite concentrations. All cultures were performed in MEM. IL-8 was quantified by ELISA after 24 hours of culture. Each point indicates the average of 4 technical replicates from a single biological replicate. Lines connect results from experiments performed on the same day. Significance was tested by unpaired one-way ANOVA followed by Dunnett's post-test with MEM + IL-1β as the reference. *$p<0.05$. C) CFTR-/- Caco-2 cells were cultured for 2 weeks in 24-well plates and then treated with IL-1β alone, or with the addition of 20mM of each metabolite unless otherwise indicated. All cultures were performed in MEM. IL-8 was quantified by ELISA after 24 hours of culture and normalized to the MEM + IL-1β control condition.

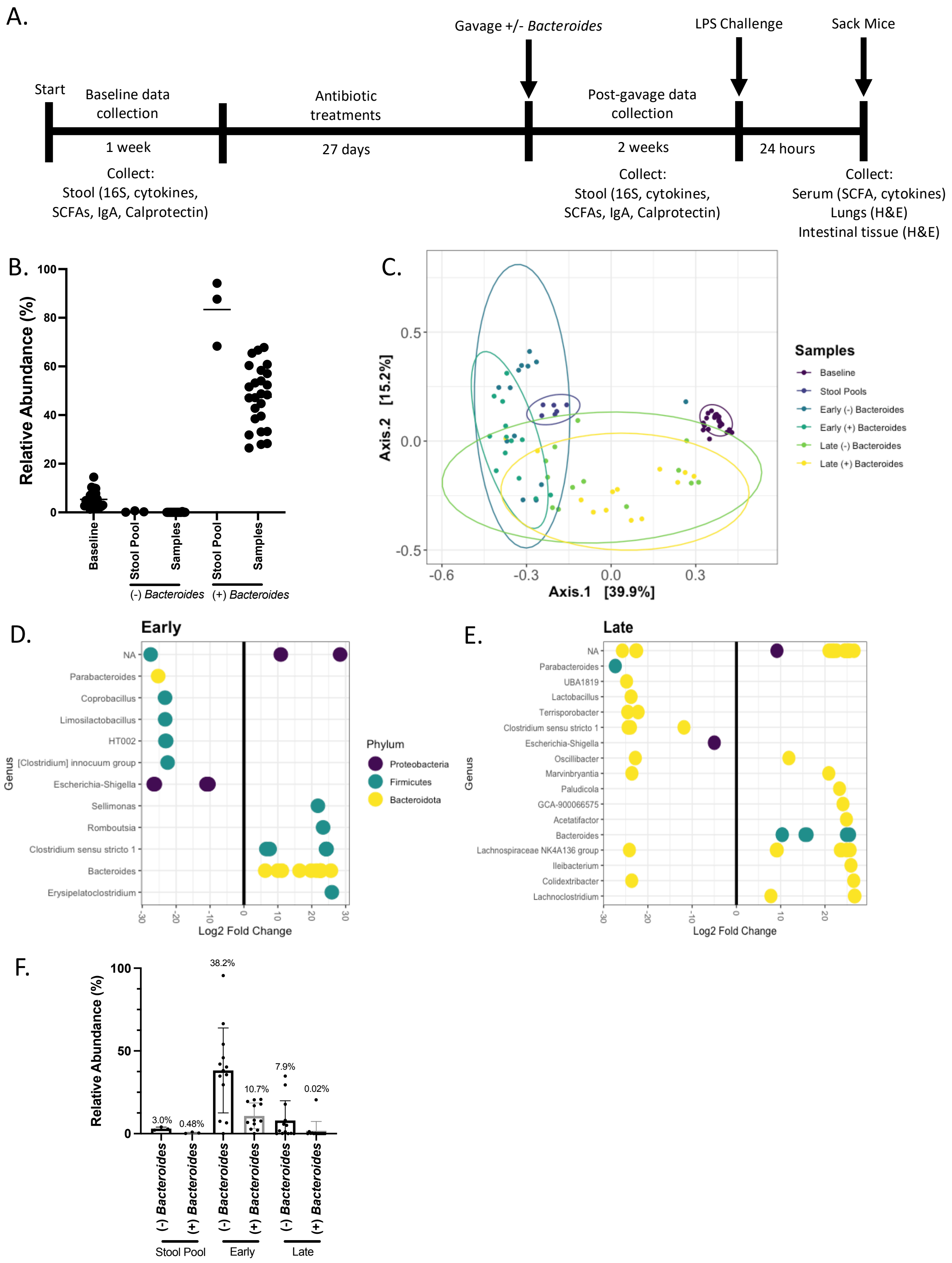
A.
Start
Baseline data collection
1 week
Collect:
Stool (16S, cytokines, SCFAs, IgA, Calprotectin)
Antibiotic treatments
27 days
Gavage +/- *Bacteroides*
Post-gavage data collection
2 weeks
Collect:
Stool (16S, cytokines, SCFAs, IgA, Calprotectin)
LPS Challenge
24 hours
Sack Mice
Collect:
Serum (SCFA, cytokines)
Lungs (H&E)
Intestinal tissue (H&E)
B.
Relative Abundance (%)
Baseline
Stool Pool
Samples
Stool Pool
Samples
(-) *Bacteroides*
(+) *Bacteroides*
C.
Axis.2 [15.2%]
Axis.1 [39.9%]
Samples
Baseline
Stool Pools
Early (-) Bacteroides
Early (+) Bacteroides
Late (-) Bacteroides
Late (+) Bacteroides
D.
Early
Genus
NA
Parabacteroides
Coprobacillus
Limosilactobacillus
HT002
[Clostridium] innocuum group
Escherichia-Shigella
Sellimonas
Romboutsia
Clostridium sensu stricto 1
Bacteroides
Erysipelatoclostridium
Log2 Fold Change
Phylum
Proteobacteria
Firmicutes
Bacteroidota
E.
Late
Genus
NA
Parabacteroides
UBA1819
Lactobacillus
Terrisporobacter
Clostridium sensu stricto 1
Escherichia-Shigella
Oscillibacter
Marvinbryantia
Paludicola
GCA-900066575
Acetatifactor
Bacteroides
Lachnospiraceae NK4A136 group
Ileibacterium
Colidextribacter
Lachnoclostridium
Log2 Fold Change
F.
Relative Abundance (%)
3.0%
0.48%
38.2%
10.7%
7.9%
0.02%
(-) *Bacteroides*
(+) *Bacteroides*
(-) *Bacteroides*
(+) *Bacteroides*
(-) *Bacteroides*
(+) *Bacteroides*
Stool Pool
Early
Late

**Figure S11. Mice treated with antibiotics and gavaged with CF stool +/- *Bacteroides*.** A) Outline of mouse experimental timeline. B) Relative abundance of *Bacteroides* in stool pools used for gavage and mouse stool collected at baseline and post-gavage from mice with the indicated condition. Each point indicates a single sample. Two stool samples (one collected early and one collected late post-gavage) are included from each mouse. C) Community structure was analyzed by multidimensional scaling (MDS) ordination based on Bray-Curtis dissimilarity coefficients and significance was analyzed by Permanova. Stool samples are grouped by Early (2-4 days) and Late (12-13 days) post-gavage collection dates. The two *Bacteroides* conditions were significantly different Early ($p = 0.014$) but not Late post-gavage ($p = 0.085$). D-E) Significant alterations in specific ASVs was analyzed by DESeq2. The Log2 Fold change of the (+) *Bacteroides* versus (-) *Bacteroides* condition is displayed on the x-axis. Each point represents and individual ASV and is labeled by genus on the y-axis and color coded by phylum. F) Relative abundance of *Eschirichia-Shigella* ASVs in stool pools used for gavage and mouse stool collected Early (2-4 days) or Late (13-13 days) post-gavage from mice with the indicated condition.

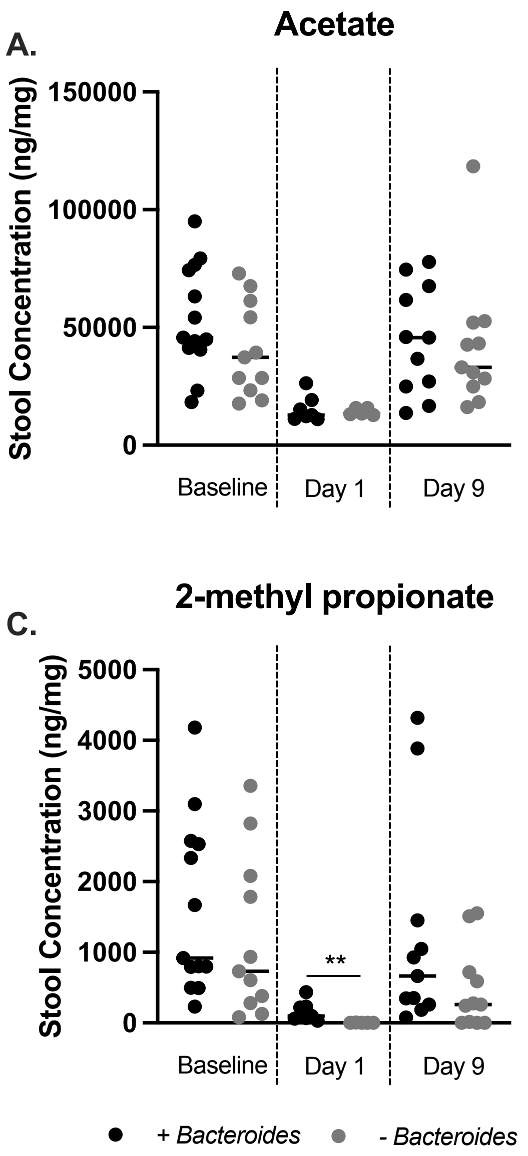


**Figure S12. SCFA quantification in mouse stool.** SCFAs A) acetate, B) butyrate, C) 2-methyl propionate, and D) 3-methyl butyrate were quantified in mouse stool by GC-MS at baseline or at the indicated number of days post-gavage. The bottom left figure legend indicates the condition for all panels. Statistical significance was analyzed by linear model between the (-) *Bacteroides* and (+) *Bacteroides* conditions for each day. All significant and marginally significant results are reported on the graph. ns = not significant. * p<0.05. ** p< 0.01.

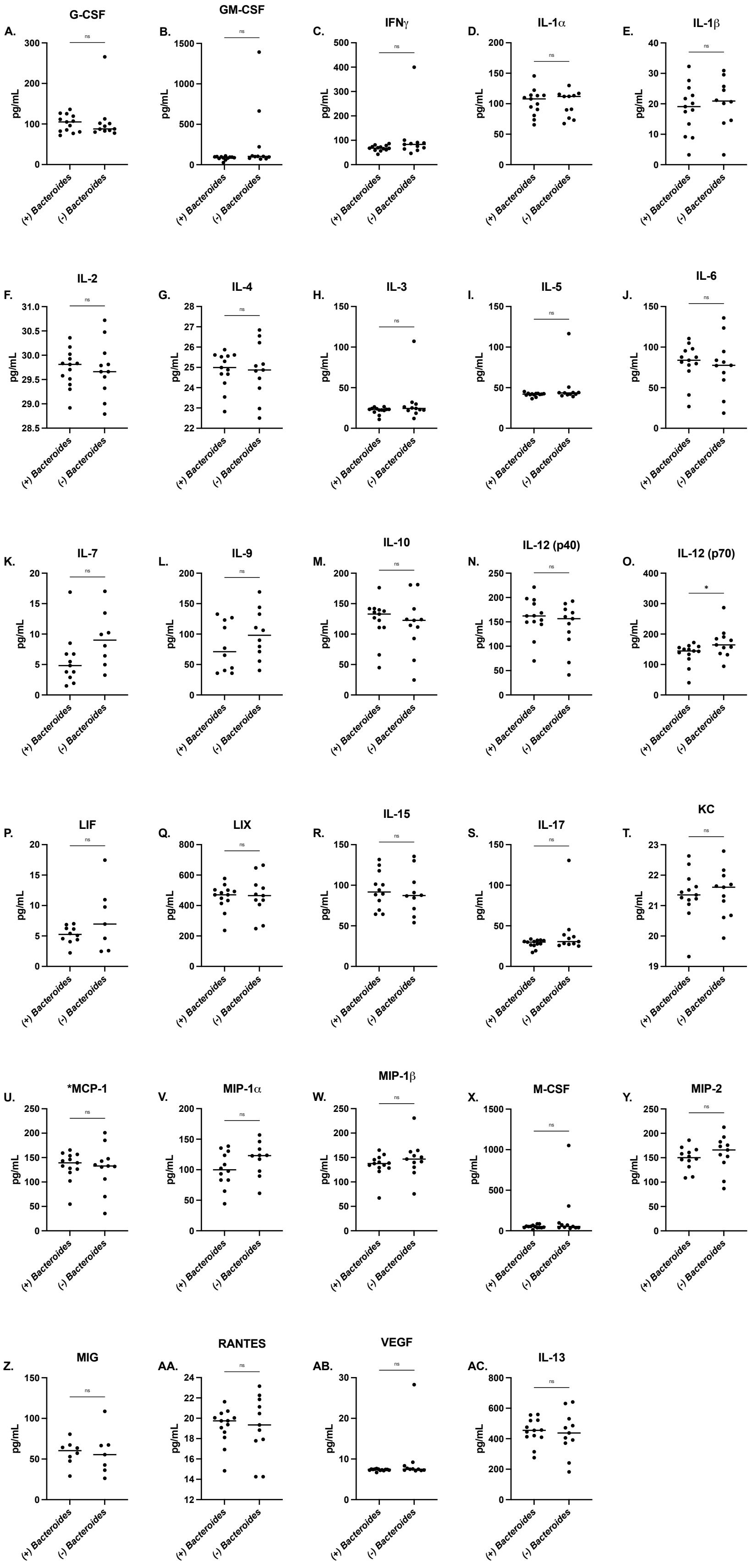

A.
G-CSF
B.
GM-CSF
C.
IFNγ
D.
IL-1α
E.
IL-1β
F.
IL-2
G.
IL-4
H.
IL-3
I.
IL-5
J.
IL-6
K.
IL-7
L.
IL-9
M.
IL-10
N.
IL-12 (p40)
O.
IL-12 (p70)
P.
LIF
Q.
LIX
R.
IL-15
S.
IL-17
T.
KC
U.
*MCP-1
V.
MIP-1α
W.
MIP-1β
X.
M-CSF
Y.
MIP-2
Z.
MIG
AA.
RANTES
AB.
VEGF
AC.
IL-13
pg/mL
(+) Bacteroides
(-) Bacteroides
ns
*

**Figure S13. Cytokines detected in mouse stool.** Mouse stool samples from three independent experiments were diluted 1:50 by weight in extraction buffer (Immunodiagnostik KR6936) and further diluted 1:5 in PBS prior to cytokine quantification by Luminex 32-plex. Each cytokine is indicated above the graph in panels A-AC. Each point indicates a single stool sample from a different mouse, and the bar indicates the mean of all samples. In the (+) *Bacteroides* condition n=13, and in the (-) *Bacteroides* condition n=11. Cytokines where fewer than 5 of the samples had cytokine within the limit of detection (LOD) (TNF$\alpha$, Eotaxin, and IP-10) were excluded from the analysis.Samples with values below the limit of detection (LOD) were excluded. Statistical significance was analyzed by linear model and pairwise results are reported in each panel. ns = not significant. * $p<0.05$.

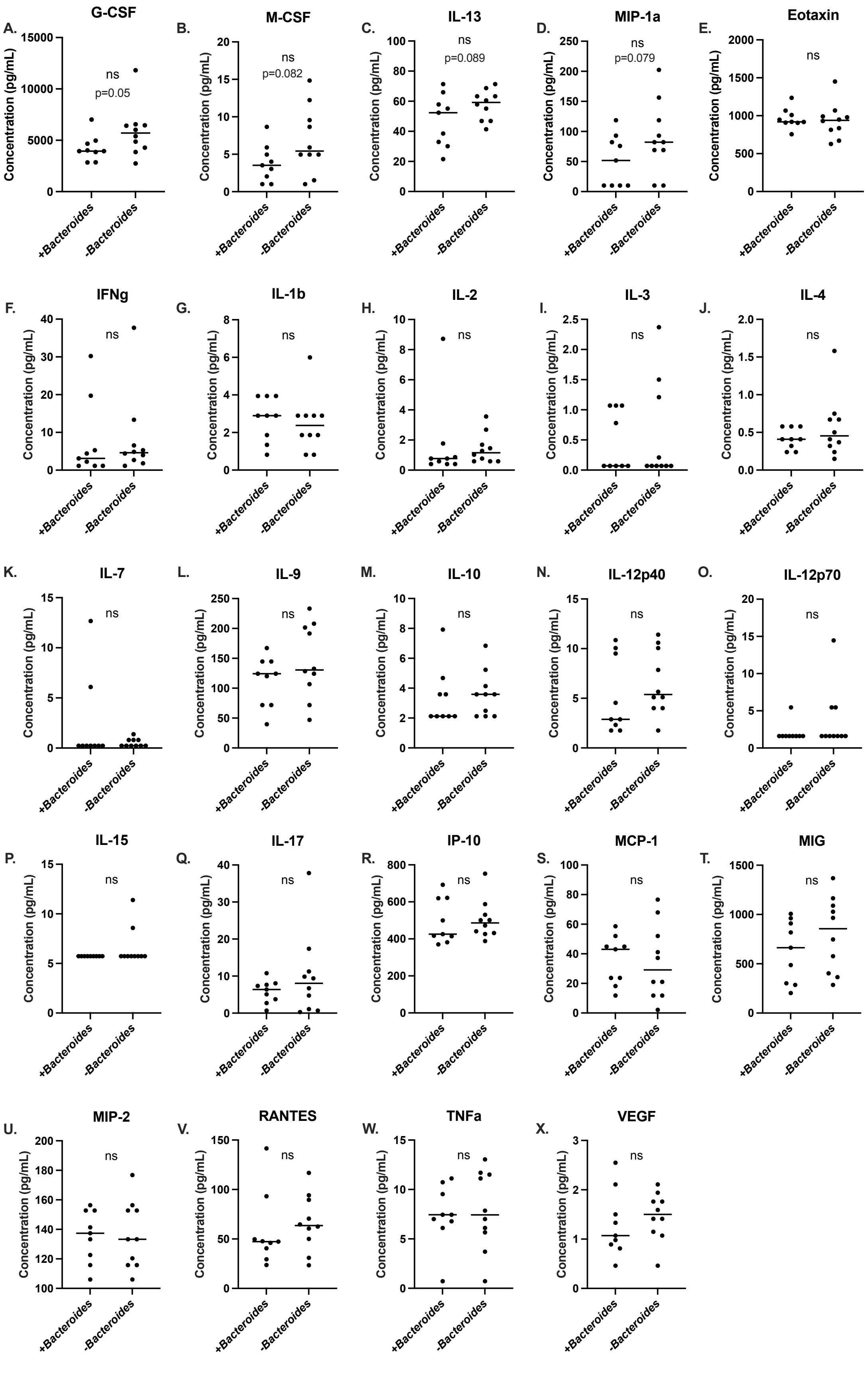
A.
G-CSF
ns
p=0.05
B.
M-CSF
ns
p=0.082
C.
IL-13
ns
p=0.089
D.
MIP-1a
ns
p=0.079
E.
Eotaxin
ns
F.
IFNg
ns
G.
IL-1b
ns
H.
IL-2
ns
I.
IL-3
ns
J.
IL-4
ns
K.
IL-7
ns
L.
IL-9
ns
M.
IL-10
ns
N.
IL-12p40
ns
O.
IL-12p70
ns
P.
IL-15
ns
Q.
IL-17
ns
R.
IP-10
ns
S.
MCP-1
ns
T.
MIG
ns
U.
MIP-2
ns
V.
RANTES
ns
W.
TNFa
ns
X.
VEGF
ns
Concentration (pg/mL)
+Bacteroides
-Bacteroides

**Figure S14. Cytokines detected in mouse serum.** Mouse serum samples collected from two independent experiments were diluted 1:2 in PBS prior to cytokine quantification by Luminex 32-plex. Each cytokine is indicated above the graph in panels A-X. Each point indicates cytokine quantity in a single serum sample from an individual mouse, and the bar indicates the mean of all samples. In the (+) *Bacteroides* condition n=9, and in the (-) *Bacteroides* condition n=10. Cytokines where fewer than 5 of the samples had cytokine within the limit of detection (LOD) (GM-CSF, LIF, and LIX) were excluded from the analysis. Samples with values below the LOD were thresholded to the minimum LOD. Statistical significance was analyzed by linear model and pairwise results are reported in each panel. ns = not significant.

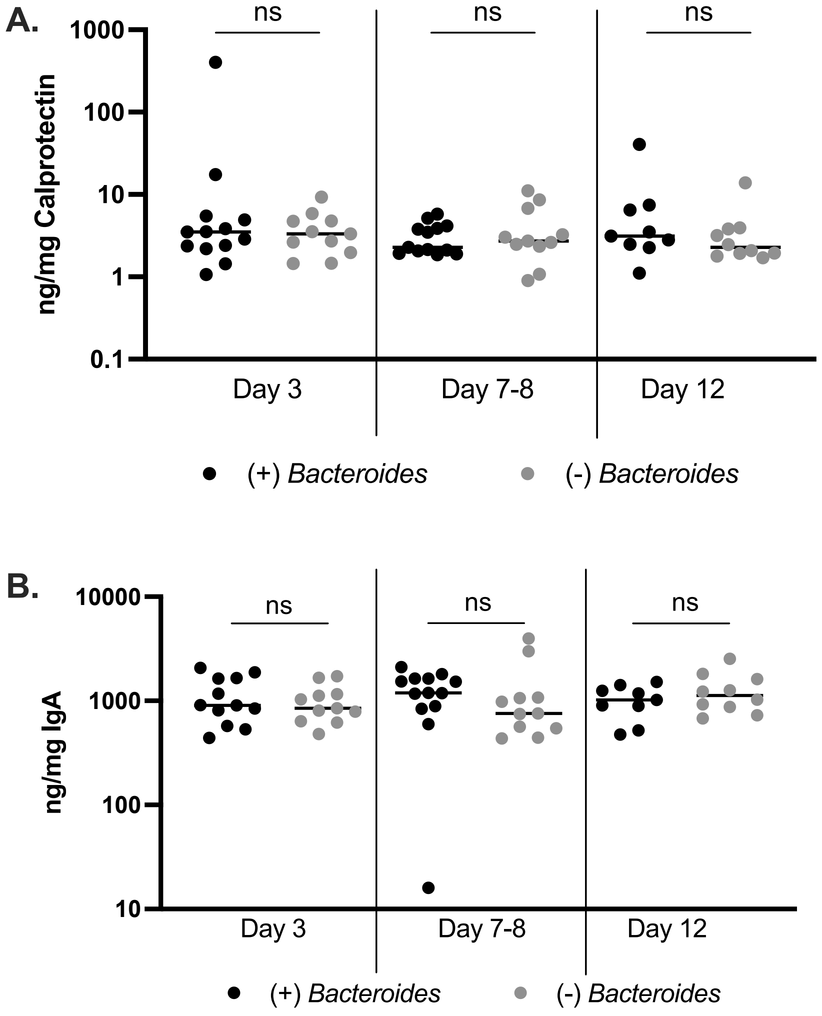


**Figure S15. Quantification of stool calprotectin and IgA.** Mouse stool samples were diluted 1:50 by weight in extraction buffer (Immunodiagnostik KR6936) and then separately quantified by ELISA for A) calprotectin and B) IgA. Statistical significance was tested by unpaired t-test for the indicated pairwise comparisons. ns = not significant. Calprotectin is a marker typically used to monitor inflammation in the gut of pwCF and other gastrointestinal disorders (14, 32, 33). Additionally, IgA, the primary immunoglobulin secreted in the intestinal tract, is increased in IBD but has not, to our knowledge, been previously assessed in pwCF or CF mice.

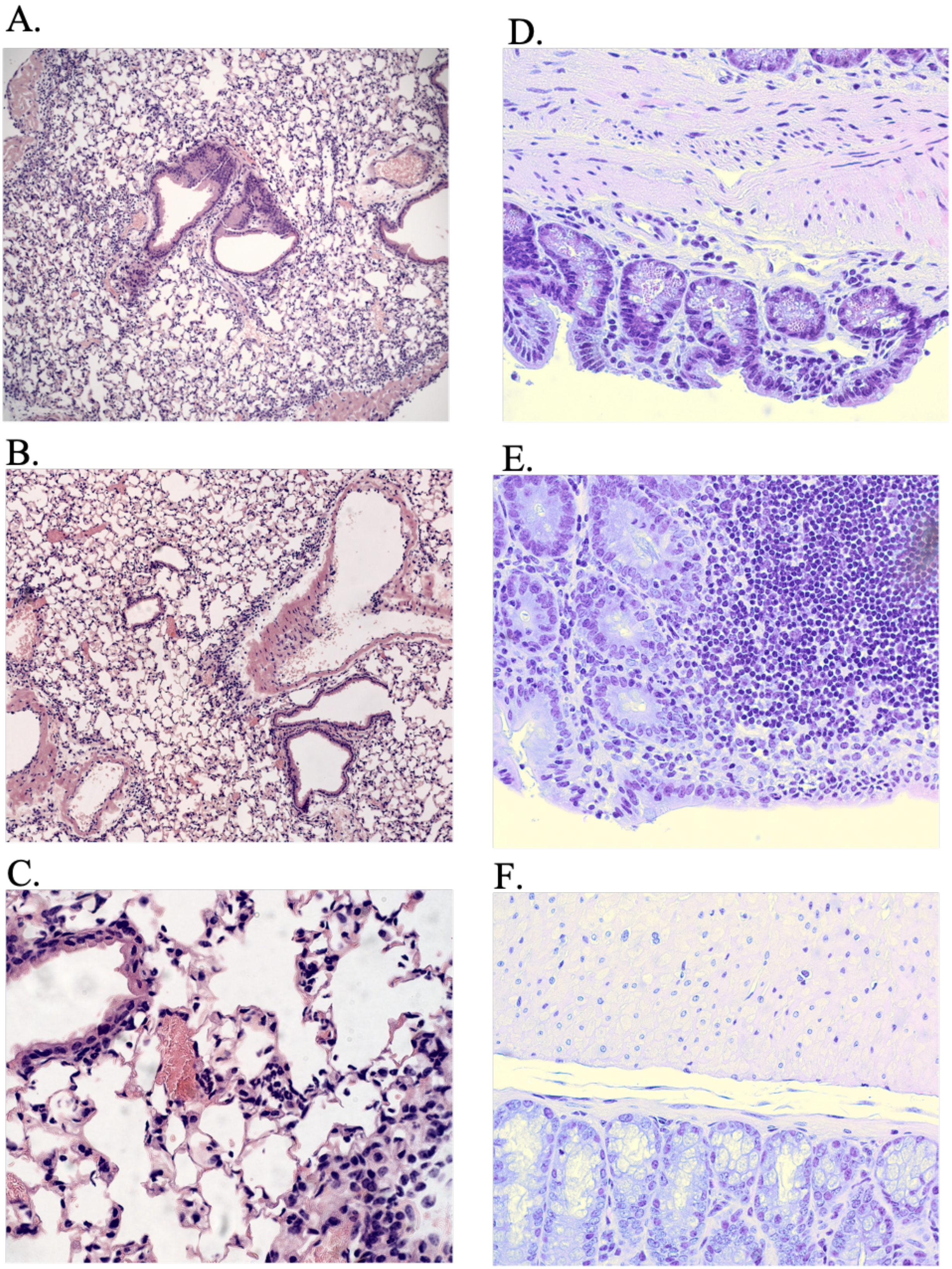
A.
B.
C.
D.
E.
F.

**Figure S16**. Lung and intestinal tissue were collected at sacrificing and preserved in 10% buffered formalin. After a minimum of 24 hours of fixation, samples were transferred to 70% ethanol and stained with Hematoxylin and Eosin (H&E) to visualize infiltration of immune cells. A-C) Representative images from lung tissue. All lung tissue demonstrated mild to no inflammatory response and there was no pattern based on intestinal *Bacteroides* condition. A) 10X image showing normal architecture without inflammatory infiltrate. B) 10X image showing a slight perivascular neutrophil infiltrate C) 40X of the same section from panel B showing a slight perivascular neutrophil infiltrate. D-F) Representative images from intestinal tissue. No significant inflammatory response was observed in any of the intestinal tissue samples. D-E) 40X images of (+) *Bacteroides* condition. Panel E includes normal gut-associated lymphoid tissue. F) 40X image of (-) *Bacteorides* condition.

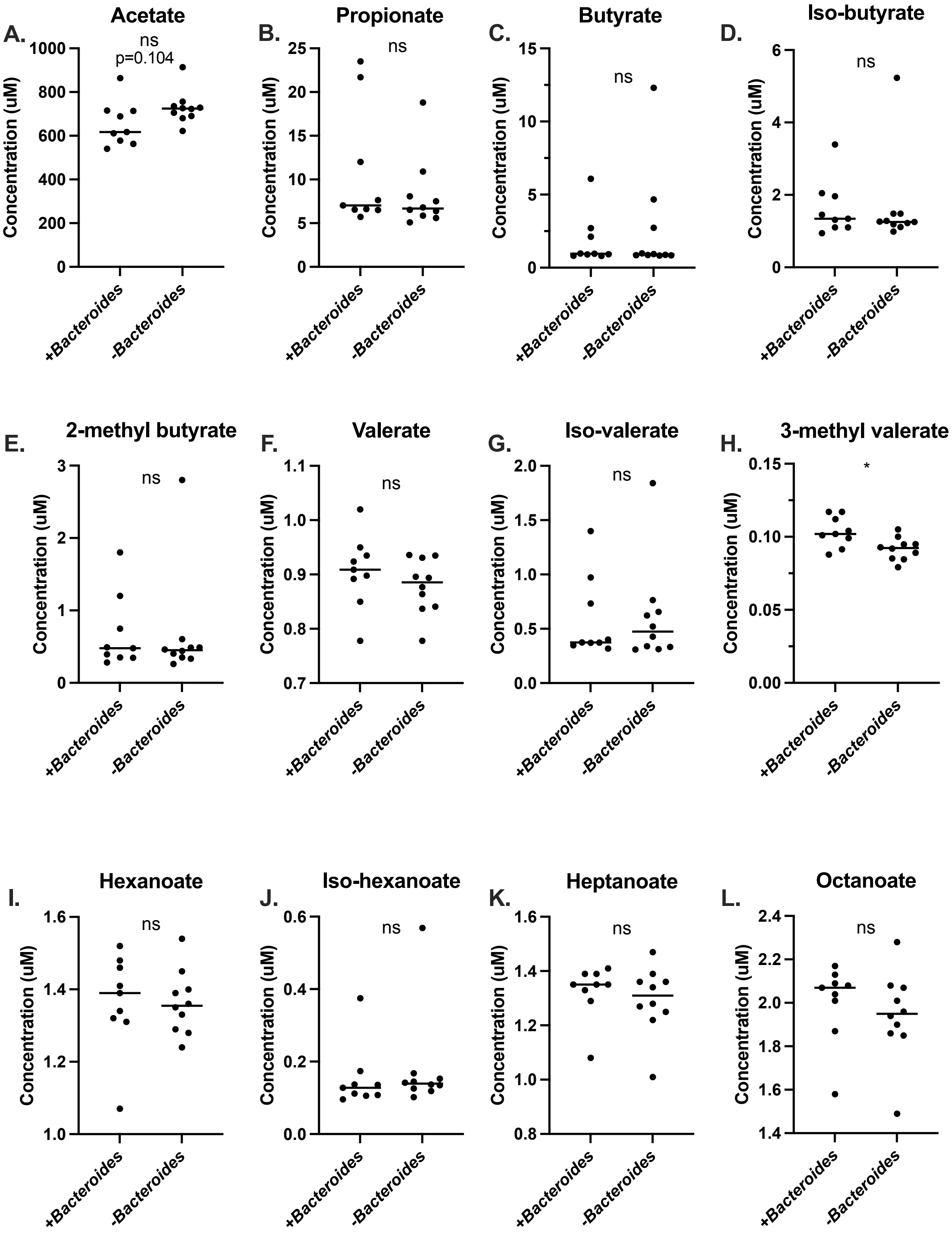

A.
Acetate
ns
p=0.104
B.
Propionate
ns
C.
Butyrate
ns
D.
Iso-butyrate
ns
E.
2-methyl butyrate
ns
F.
Valerate
ns
G.
Iso-valerate
ns
H.
3-methyl valerate
*
I.
Hexanoate
ns
J.
Iso-hexanoate
ns
K.
Heptanoate
ns
L.
Octanoate
ns
Concentration (uM)
+Bacteroides
-Bacteroides

**Figure S17. SCFA quantification in mouse serum.** SCFAs were quantified in mouse serum by LC-MS/MS (Duke Proteomics and Metabolomics Shared Resource). Each SCFA is indicated above the graph in panels A-L. Statistical significance was analyzed by linear model and pairwise results are reported in each panel. ns = not significant. * $p<0.05$.

Hao, S., Roesch, E. A., Perez, A., Weiner, R. L., Henderson, L. C., Cummings, L., . . . Drumm, M. L. (2020). Inactivation of CFTR by CRISPR/Cas9 alters transcriptional regulation of inflammatory pathways and other networks. *J Cyst Fibros, 19*(1), 34-39. doi:10.1016/j.jcf.2019.05.003
